# Cell-specific RNA isoform remodeling in the aging mouse brain

**DOI:** 10.1101/2025.06.05.658133

**Authors:** Abid Rehman, Megan Duffy, Katarina Gresova, Cedric Belair, Na Yang, Lin Wang, Christopher T. Lee, Matthew Payea, Sulochan Malla, Showkat Ahmad Dar, Payel Sen, Mark R. Cookson, Manolis Maragkakis

## Abstract

Aging is associated with increases in risk of multiple chronic diseases and at the cellular level, with disruptions in RNA homeostasis. Single-cell transcriptomic studies have revealed RNA abundance heterogeneity across brain cell types but the more complex changes in RNA processing such as alternative splicing, isoform usage, transcription start and polyadenylation site selection remain uncharacterized. Here, we combine single-cell analysis with long-read nanopore sequencing to capture full-length RNA isoforms and uncover temporal changes in RNA transcription, processing, and alternative splicing in the aging mouse cortex and hippocampus. By profiling transcriptomes from young adult to very old mice, we identify non-linear, cell-type-specific isoform expression changes and isoform usage shifts, primarily driven by transcription start site selection. These aging-associated isoform changes alter the coding potential and poly(A) site position of genes. Our data also reveal a high proportion of senescence in immune cells, far exceeding that of other cell types. We also identify isoform markers that, when applied to a machine learning model, distinguish senescent from normal immune cells. This study provides a full-length RNA isoform-based atlas of the aging mouse brain, offering insights into RNA metabolism remodeling across brain cell types throughout the lifespan.

## Introduction

Brain aging is associated with alterations in physiological and biochemical processes that increase the risk of developing disease including neurodegenerative disorders ^1^. Development of novel approaches to mitigate aging effects could have great promise for disease prevention and cognition maintenance but will require discovery of the underlying temporal molecular changes^2^. The mammalian brain comprises cells of several distinct types with high spatial and cellular heterogeneity thus making it challenging to identify cell-type specific molecular changes across lifespan ^3–5^.

Single-cell transcriptomics has revealed a strong link between aging and transcriptomic changes ^3,6–11^. Following these and other studies ^12–15^, deregulation of the transcriptome has recently been proposed as a central feature of aging, although relatively little is known about the molecular events involved ^16–18^. Typically, single-cell transcriptomic studies are limited to measurements of steady-state RNA abundance without the ability to further explore the underlying changes in transcription, splicing, processing, and decay ^19,20^. Therefore, much less is known about the cell-type specific regulation of these defining components of RNA metabolism in the adult and aging mammalian brain.

A major limitation to investigating changes in RNA metabolism has been the reliance on short-read sequencing. However, long-read sequencing has recently been used to capture transcriptome diversity across different tissues and ages ^21,22^, as well as identify novel modes of RNA decay and isoform usage ^23–25^, thus addressing important aspects of RNA metabolism. Combining long-read with single-cell sequencing has therefore provided important insights on previously uncharacterized transcriptomic patterns during development of the mouse and human brain ^26,27^. However, a single-cell map of RNA isoforms to allow characterization of the changes and the cellular heterogeneity of RNA metabolism in the aging brain is currently unavailable.

Here we combine single-cell and long-read sequencing to discover changes in RNA metabolism in the aging brain, specifically of transcription start site (TSS) selection, alternative splicing, and transcription termination. We measure heterogeneity across the brain by independently profiling two regions critical to aging biology, the cortex and hippocampus. Importantly, we identify sex differences and non-linear trends that span young, middle-age, old and geriatric animals. Our results reveal distinct and non-linear RNA isoform expression trajectories and switches across age, cell type, and sex. Isoform changes across time are dominated by TSS changes and select against non-coding RNA expression. Our data also reveal that immune cells have much higher proportions of senescent cells than other cell types. We further identify isoform markers that, used in a machine learning model, can distinguish senescent from normal immune cells. Collectively, our work provides an RNA isoform atlas of single cells in the aging brain and reveals dynamic changes of RNA metabolism across lifespan.

## Results

### Mapping the cellular and regional transcriptomic landscape of the aging mouse brain with long-read sequencing

To uncover the lifetime changes and the cellular heterogeneity of the mammalian brain with respect to regulating RNA processing and metabolism, we combined 10X single-cell library preparation with Oxford Nanopore Technology long-read nanopore sequencing. To capture the progressive nature of transcriptomic changes across time, we separately collected samples from four timepoints: young (4 months), middle (16 months), old (22-23 months), and geriatric (27-31 months) animals. Additionally, we independently profiled the cortex and hippocampus of both male and female mice for a total of 51 processed samples (**Fig. 1A, Sup. Table 1**).

**Fig. 1:**
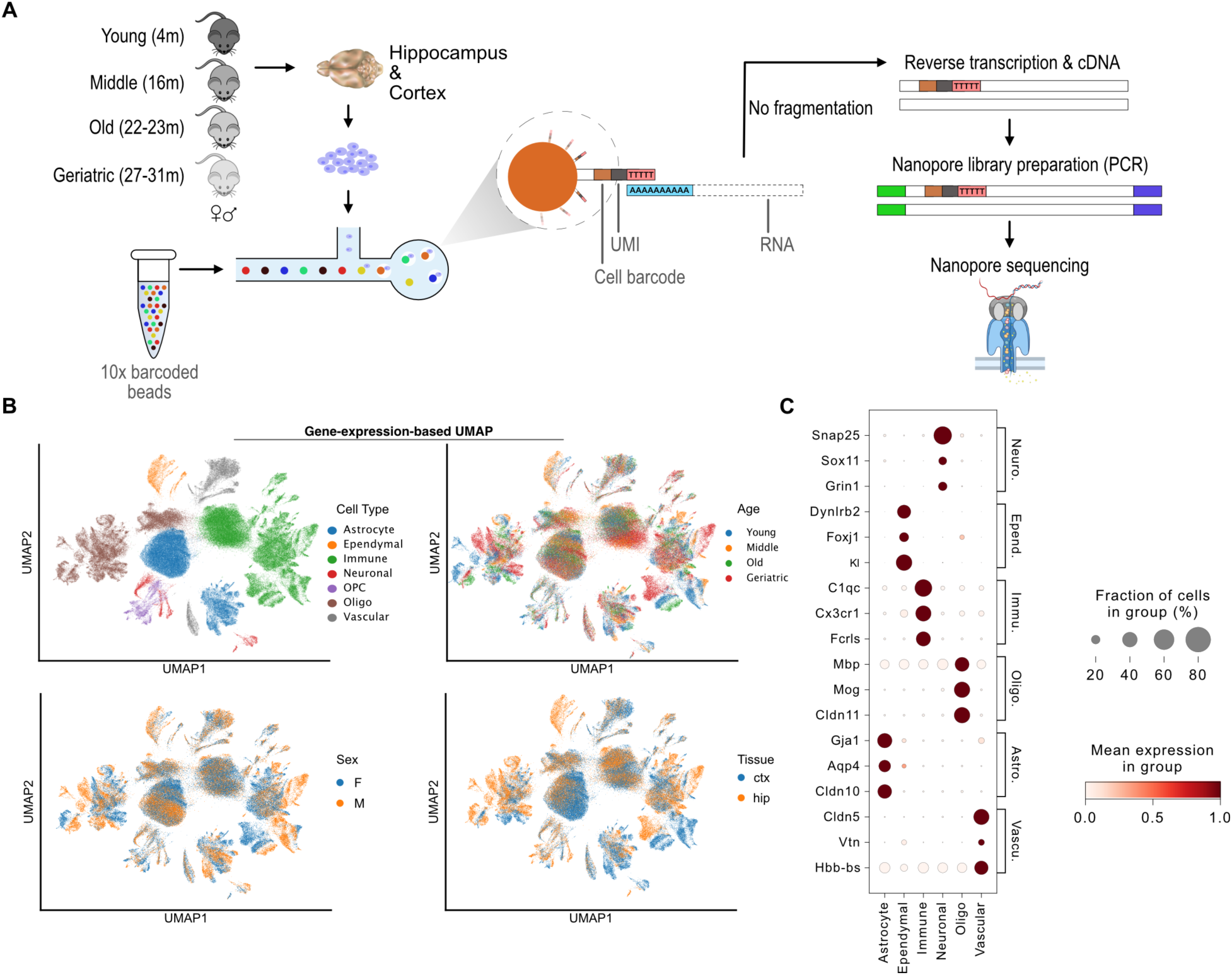
Mapping the cellular and regional transcriptomic landscape of the aging mouse brain with long-read sequencing. **A)** Schematic of experimental workflow. **B)** Scatter plots of UMAP gene abundance embeddings of all cells. Each point represents a cell and is colored by the identified cell type or the corresponding sample age, sex and brain region. **C)** Dot plot of marker gene expression across cell types.

To enrich for mature RNAs over molecules containing unspliced intronic sequences, we extensively optimized single cell rather than single nuclei isolation. Following sample isolation, cell dissociation, and 10X droplet formation, RNA libraries from non-fragmented RNA were prepared and sequenced on a Nanopore PromethION device (**Fig. 1A**). Overall, approximately 45 terabytes of raw nanopore data were processed comprising over 10 billion long reads (**Sup. Fig. 1A**) that resulted in an average of 3,077 cells per sample. Cells with mitochondrial content above 20%, likely dying cells, or fewer than 100 quantified genes (**Sup. Fig. 1B, C**) were discarded to ensure substantial depth for subsequent gene and isoform expression analysis. Following this filtering, 156,935 cells were retained for further analysis.

While our long-read sequencing data directly quantified individual RNA isoforms, we initially aggregated the data at gene level to facilitate cell-type annotation. We integrated all cells across all samples and then assigned cell types to each computationally identified cell cluster using marker genes from the PanglaoDB ^28^ and Tabula Muris ^29^ databases (**Fig. 1B-C, Sup. Fig. 2-3**). In total, seven cell types were identified: neurons, oligodendrocytes, oligodendrocyte precursor cells (OPCs), astrocytes, ependymal, vascular, and immune (**Fig. 1B, Sup. Table 2**). Ependymal cells were primarily derived from a single sample and thus were excluded from downstream analysis while OPCs, due to their low numbers, were incorporated into the broad oligodendrocyte lineage (**Sup. Fig. 4A**). Quantification of cell type composition showed a similar distribution between males and females **(Sup. Fig. 4B)** while, as expected ^30^, substantial differences were found between brain regions, with the cortex containing significantly more astrocytes than the hippocampus **(Sup. Fig. 4C)**. Overall, we found that the single-cell brain data primarily clustered by cell type while also displaying separation by sex, brain region, and age.

### Transcription start site selection drives isoform specificity across cell types

Having established a cellular map based on gene abundance, we subsequently enhanced the granularity of the projection by incorporating RNA isoform abundance as quantified by long read sequencing. To achieve this, we clustered cells based on isoform abundance and subsequently projected the already established cell-type annotations on the isoform cellular map (**Fig. 2A**). Analysis of cell type composition in the isoform-defined cell clusters showed high cellular specificity in distinct clusters indicating robust transcriptomic separation (**Sup. Fig. 5A**).

**Fig. 2:**
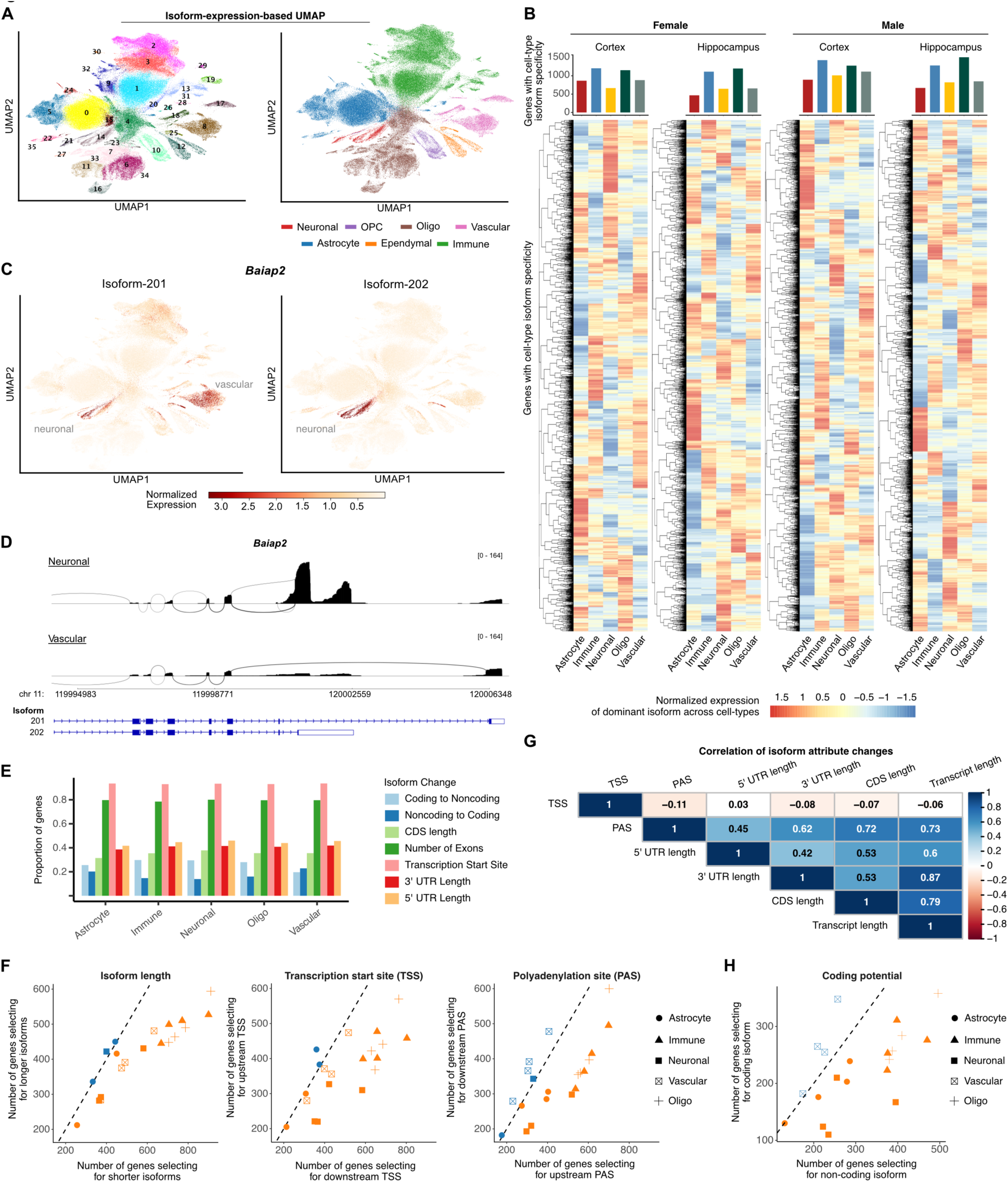
Transcription start site selection drives isoform specificity across cell types. **A)** Scatter plot of UMAP isoform expression embeddings of all cells colored by Leiden-identified clusters (left) and cell type (right). **B)** Heatmap of isoform expression of genes with significant cell-type isoform specificity in indicated cell-type compared to other cell types. The number of genes with significant isoform specificity in each cell-type is shown in bar plots at the top. Clustering along the y-axis is performed independently for each sex, brain region combination **C)** Scatter plot of UMAP isoform expression embeddings with cells colored by the expression of Baiap2 isoforms, 201 and 202. **D)** Sashimi plots of gene Baiap2 in the male cortex, comparing isoform expression between neuronal and vascular cells. **E)** Bar plot of proportion of significant isoform-specific genes stratified by the type of change between the cell-specific and non-specific isoform. **F)** Scatter plot of the number of genes where cell-type-specific isoforms are shorter/longer, use upstream/downstream TSSs and upstream/downstream PASs, compared to non-specific ones. For each cell type, the four sex and brain region combinations are shown as independent points. **G)** Spearman’s correlation coefficients of isoform attribute changes for CIGs with at least two protein-coding transcripts. For example, PAS is correlated with transcript length indicating that downstream PAS is associated with longer isoforms. **H)** Scatter plot of the number of genes where cell-type-specific isoforms are non-coding, compared to non-specific ones. For each cell type, the four sex and brain region combinations are shown as independent points.

To explore isoform heterogeneity across brain cell types, we searched for genes ubiquitously expressed across most cell types, but which expressed distinct, cell-type specific RNA isoforms. We applied Dirichlet-multinomial modeling of RNA isoform usage ^31^ comparing each cell type with all others. In total, we identified 3,769 Cell-type Isoform-specific Genes (CIGs) (p < 0.05, **Fig. 2B, Sup. Table 3-4)**. Out of all CIGs, 67% were shared between sexes and 68.5% between the cortex and hippocampus revealing a high overlap between biological factors (**Sup. Fig. 5B**). Overall, we found that astrocytes and neurons consistently had the lowest number of CIGs across both sexes and brain regions. In contrast, immune cells and oligodendrocytes had the highest number of CIGs, indicating they expressed a more selective transcriptomic program, possibly the result of higher underlying cell subtype heterogeneity (**Fig. 2B**). An interesting example of a CIG is *Baiap2*, a brain-specific angiogenesis inhibitor (BAI1)-binding protein. *Baiap2* expressed two distinct isoforms: isoform 201, found in both vascular and neuronal cells, and isoform 202, exclusively expressed in neuronal cells (**Fig. 2C**). These two isoforms are annotated with distinct terminal coding and 3’ untranslated regions (3’ UTRs) indicating a unique regulatory profile in the two cell types. Specifically, isoform 202, expressed in neuronal cells, includes a longer, upstream 3’ UTR compared to isoform 201, expressed in vascular cells (**Fig. 2D**).

Gene isoforms can differ in multiple attributes, including TSS, exon and intron composition, coding sequence, and UTR length. Additionally, some isoforms of coding genes may lack coding potential, conferring a regulatory role via the nonsense-mediated decay pathway ^32^. To assess isoform attribute changes, we analyzed CIGs by selecting the two isoforms per CIG with the greatest increase and decrease in usage relative to other cell types, comparing 3,769 isoform pairs. Our analysis identified TSS (92%) and exon count (75%) as the most frequently altered attributes across sexes and brain regions (**Fig. 2E, Sup. Fig. 5C**). However, considering only two isoforms per CIG may overlook simultaneous changes in multiple isoforms of a gene. To address this, we developed a weighted attribute metric, incorporating all isoforms of a gene. Specifically, we weighted each attribute (e.g., length, TSS, poly(A) site, and coding potential) by its relative isoform usage, defined as the proportion of gene expression attributed to an isoform. Using this approach, our data showed that CIGs in all cell-types, except astrocytes, displayed specificity towards increased usage of shorter isoforms compared to other cell types (**Fig. 2F**). This pattern likely suggests an evolutionary selection process in which cell-type-specific isoforms arose as variants from within existing gene boundaries rather than intergenic sequence space. Consistent with this, our results also showed increased usage of downstream TSS and upstream poly(A) sites, particularly for immune, neuronal and oligodendrocyte cells (**Fig. 2F**). *Baiap2* is an example where the neuron-specific isoform 202, uses an upstream poly(A) site compared to the non-neuron-specific 201 (**Fig. 2D**).

Recent studies in the fly and human nervous system showed that changes in TSS are consistently linked to corresponding shifts in 3′ end site selection ^33^. However, it remained unclear whether these changes were correlative such that a downstream change in the TSS would be accompanied by a downstream change in the 3′ end. Instead, we found that usage of upstream or downstream TSSs was not directly correlated with the corresponding isoform length or poly(A) site position (**Fig. 2G**). However, our data did show a high correlation between poly(A) site and isoform length (ρ = 0.73, **Fig. 2G**) indicating that selection of upstream poly(A) sites was associated with shorter isoforms.

Combined, our results identify thousands of isoforms, typically of shorter length and with downstream TSSs and upstream poly(A) sites, with selective expression in distinct cell types of the adult mouse brain.

### Brain isoform cell-type specificity is biased towards non-coding isoforms

We then focused our analysis on genes with both coding and non-coding isoforms to examine how isoform usage differences between cell types affect gene coding potential. Interestingly, we found that in all cell types, except vascular, there was higher specificity towards non-coding isoforms (**Fig. 2H**). Given that isoforms that do not code for proteins are expected to have regulatory roles on gene expression via RNA decay pathways ^32,34^ our findings indicate that isoform cell-type specificity may facilitate RNA regulation besides coding for alternative proteins. Interestingly, our data also showed positive correlation (ρ = 0.45) between poly(A) site position and coding potential, indicating that non-coding isoforms specific to a cell-type are also more likely to have upstream poly(A) sites compared to coding counterparts (**Sup. Fig. 5D**). This is consistent with our finding of CIGs preferentially using isoforms with upstream poly(A) sites (**Fig. 2F**). Nevertheless, while most cell types display specificity for non-coding isoforms and upstream poly(A) sites, vascular cells were noticeably an exception as they exhibited a modest preference for coding isoforms (**Fig. 2F**) and downstream poly(A) sites (**Sup. Fig. 5E**), highlighting a distinction between different cell-types of the brain.

Analyzing protein-coding isoforms, we also found a preference for shorter lengths across the 5ʹ UTR, coding sequence (CDS) and 3ʹ UTR (**Sup. Fig 5E**) with positive correlations between each pair of these regions (**Fig. 2G**). Further analysis revealed that over 70% of CIGs that selected for longer CDS likewise had a longer 3ʹ and 5ʹ UTR (**Sup. Fig. 5F**). We wondered whether this correlation between change in the 5ʹ UTR, CDS and 3ʹ UTR was to be expected due to background distribution of the annotated transcriptome. We thus quantified the corresponding correlations in the annotated transcriptome without factoring in the weighted usage values. Our results found no correlation between these attributes in the annotated transcriptome (**Sup. Fig. 5G**). These results indicated that cell-type isoform specificity involved coupled, concomitant changes in the CDS, 5ʹ and 3ʹ UTR. Collectively, our data suggest that cell-type isoform specificity regulates coding potential with a preference for non-coding isoforms, and an associated selection of upstream poly(A) sites, or coding isoforms with shorter genic features including the 5’ UTR, CDS and 3’UTR.

### Immune signaling is a component of isoform regulation across ages and brain cell types

These analyses identified a cell-type specific isoform expression program in distinct cell types of the mouse brain. PCA showed substantial variance with age, and we therefore wondered how this program may be changing across time (**Sup. Fig. 6**). To capture both linear and non-linear trends of isoform expression across time, we employed spline modeling coupled with likelihood ratio test to determine significant departures from a stable, flat expression trajectory. Across cell types, sexes and brain regions, our analysis identified 7189 isoforms with significant temporal deviation from a flat line with most changes observed for females and the hippocampus compared to males and cortex (**Fig. 3A**).

**Fig. 3:**
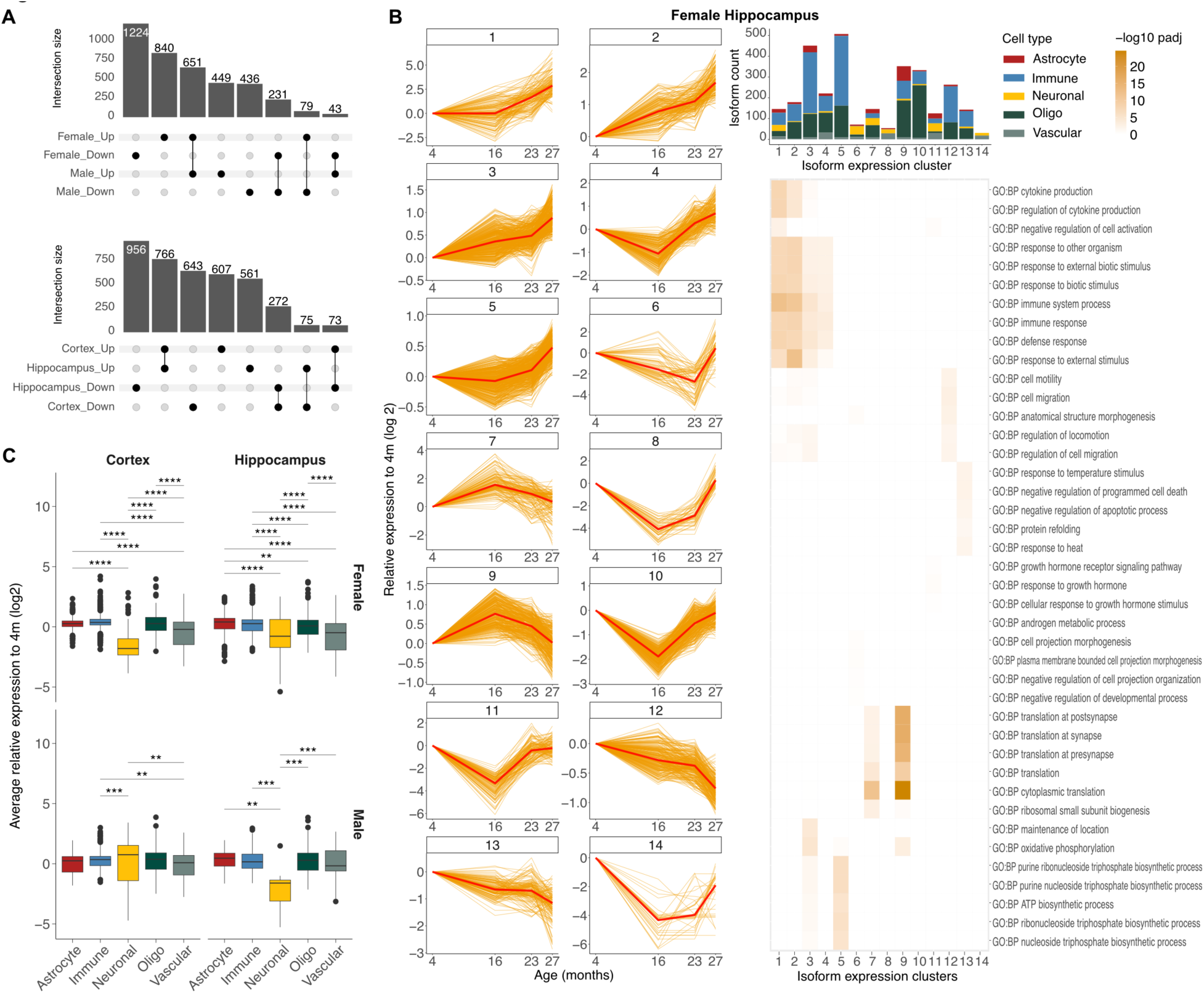
Immune signaling is a component of isoform regulation across age and brain cell types. **A)** Upset plots of the number isoforms with significant aging-associated expression change that overlap between sexes and brain regions. **B)** Line plots (left) of isoform expression at different ages in the female cortex clustered by expression trajectory. Each line represents isoform expression within a cell type, normalized to the corresponding expression at 4 months. The dark red line corresponds to the average expression of the cluster. Bar plot (top right) of cellular composition of each isoform expression cluster. Heatmap (bottom right) of significance of gene ontology terms enrichment in each cluster. Enrichment is calculated against all expressed genes in the female hippocampus. **C)** Box plot of average relative expression difference from young for each cell type.

To identify shared expression trajectories within these isoforms, we used a method employing mixed-effects models with Gaussian variables to cluster expression traces displaying similar temporal changes ^35^. To select the optimal number of temporal clusters, we compared clustering silhouette scores, quantifying inter- and intra-cluster distances, for 8, 10, 12 and 14 clusters and selected the number that maximized the score for each sex and brain region independently. Our results identified 14 temporal clusters for female hippocampus, 8 for female cortex and 10 for each of the male hippocampus and cortex (**Sup. Table 5**). We subsequently arranged the identified clusters by increasing expression fold-change of the geriatric over young age (**Sup. Table 6**). Overall, our analysis identified clusters with non-linear expression trajectories across time indicating that isoform expression changes start early in life and are dynamically regulated across the lifespan (**Fig. 3B, Sup. Fig. 7, 8A**), consistent with recent reports ^36^.

Also consistent with previous work on gene expression ^37^, analysis of isoform expression showed a modest age-associated decrease in expression of longer isoforms for all cell types, except neurons that displayed a more heterogeneous profile across sex and brain regions (**Sup. Fig. 9A**). Interestingly, we also noticed that neuronal isoforms often resided in clusters separate from other cell types and thus wondered whether a distinct aging neuronal isoform expression pattern existed (**Fig. 3A, Sup. Fig. 6A, 7A, B – bar plots**). Indeed, quantification of the average isoform expression change from middle to geriatric age compared to young, revealed that neuronal isoforms exhibited the highest aging-associated expression reduction (**Fig. 3C**). Overall, our data showed the highest overlap in expression changes between males and females as well as between the immune, oligo and astrocyte lineage. In contrast, neuronal and vascular isoforms showed the lowest overlap with other cell types (**Sup. Fig. 8B**).

Gene ontology analysis of each temporal expression cluster identified terms associated with stress and immune response as enriched in upward expression trends with age and shared across sex and brain regions (**Fig. 3B, Sup. Fig. 7, 8A - heatmaps**). Immune cells showed the strongest signal for immune response terms followed by astrocyte, oligodendrocytes and vascular, while neurons showed the lowest (**Sup. Fig. 9B**). These results are consistent with an integrated increase in inflammatory signaling across diverse cell types as previously reported in aging ^38^. Terms related to translation at synapse and presynapse were also found to be significantly altered with age across sex and brain regions, with the corresponding genes being upregulated with age. Overall, our results identify that aging is associated with widespread isoform expression remodeling across cell types of the brain. Furthermore, isoform remodeling strongly affects the neuronal transcriptome that exhibits the highest age-related downregulation.

### Isoform usage changes are prevalent in aging

Our analysis identified thousands of age-related isoform expression changes and thus we also wished to investigate whether isoform usage within genes could also change over time favoring specific isoforms over others. To explore this, we again applied a Dirichlet-multinomial model to identify Age Isoform-specific Genes (AIG) in pairwise comparisons across young, middle, old, and geriatric age groups. This analysis identified 4,717 unique AIGs in at least one comparison (**Fig. 4A, Sup. Fig. 11A, Sup. Table 7, 8**). Example AIGs for the female hippocampus are shown in **Fig. 4B** demonstrating a mix of different patterns of usage change, including progressive switch from one to another isoform or selective expression of alternative isoforms with age.

**Fig. 4:**
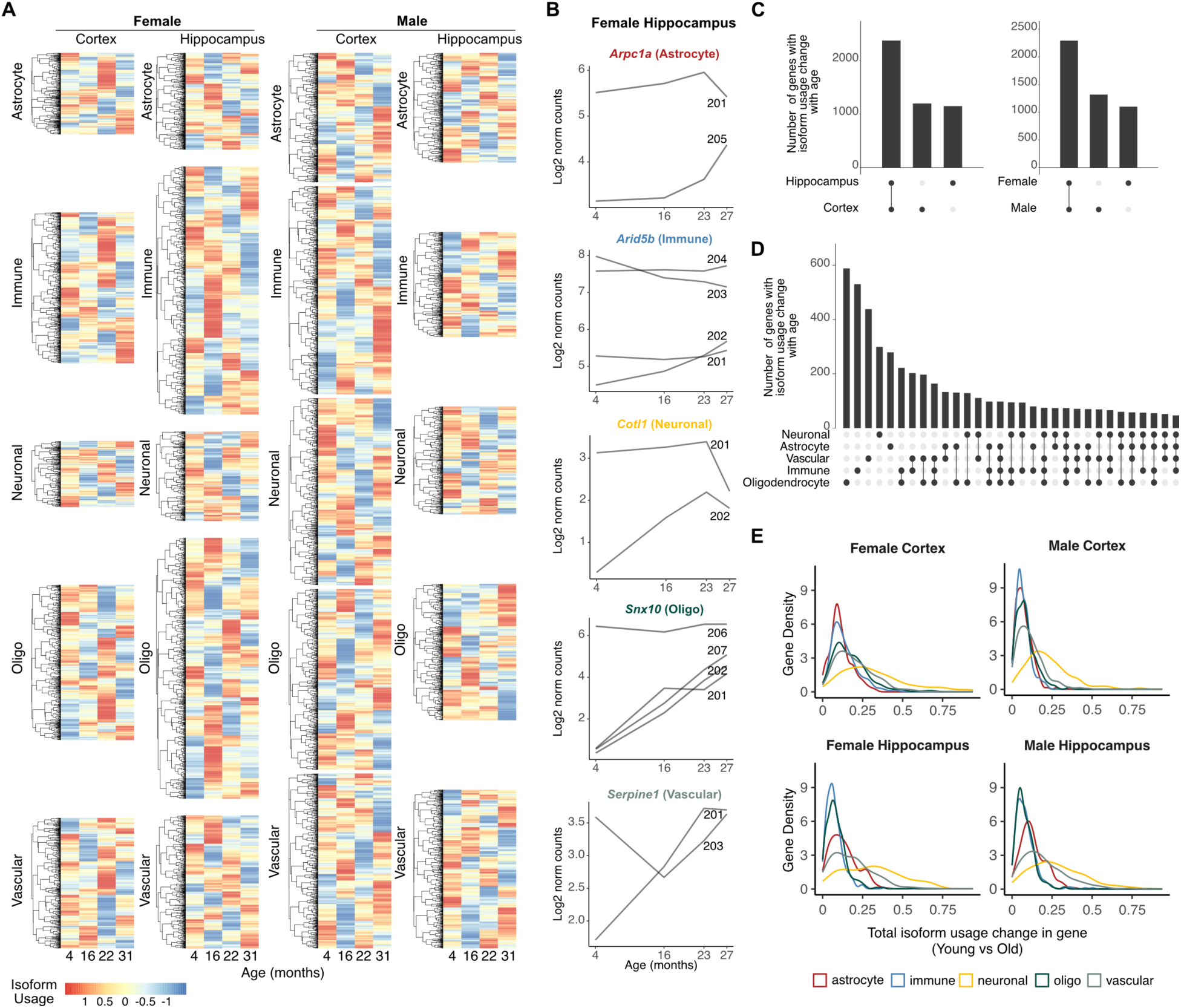
Isoform usage switching is prevalent in aging. **A)** Heatmaps of row-normalized isoform usage values across time. The usage of the most dominant isoform at young age is shown. **B)** Example line plots of isoform expression for genes with significant isoform usage changes in the female hippocampus. **C-D)** Upset plots of the number of unique and shared genes with significant isoform usage change across time between brain regions and sex (C) and cell types (D). **E)** Isoform distribution of total change in isoform usage between young and old.

Approximately half of AIGs involved only two changing isoforms, while the remaining involved three or more isoforms (**Sup. Fig. 11B**). AIGs exhibited substantial overlap across sex and brain regions, 48.7% and 50.3%, respectively (**Fig. 4C**) but greater heterogeneity was observed between individual cell types, indicating a cell-type specific isoform usage regulatory program with age (**Fig. 4D**). Specifically, oligodendrocytes demonstrated the highest overlap with other cell types, particularly with immune and vascular cells.

Interestingly, while oligodendrocyte and immune cells had numerically the highest number of AIGs (**Fig. 4A, Sup. Fig. 11A**), vascular and neuronal cells had the highest magnitude of isoform usage change (**Fig. 4E**). Gene ontology analysis of AIGs did not reveal significant enrichment indicating that likely isoform changes were not restricted to a specific biological component or function. Overall, our findings identify a widespread program of cell-type specific isoform remodeling with age.

### Isoform usage with age associates with upstream transcription and downstream poly(A) site selection

We next aimed to identify the changes in isoform attributes that were associated with the age-related usage remodeling. Similar to cell-type specificity, TSS selection and exon number were again the most dominant age-related changes, with over 89% and 79% of events involving changes in these isoform attributes, respectively (**Fig. 5A**). These results were consistent across both sexes and brain regions (**Sup. Fig. 12A**). However, in contrast to cell-type specificity, we observed an age-related shift towards selection of upstream TSS and downstream PAS, especially for immune cells (**Fig. 5B**). Similarly, a modest shift towards longer isoforms was also observed (**Sup. Fig. 12B**). These changes manifested from middle to geriatric age indicating that they initiate early in adulthood.

**Fig. 5:**
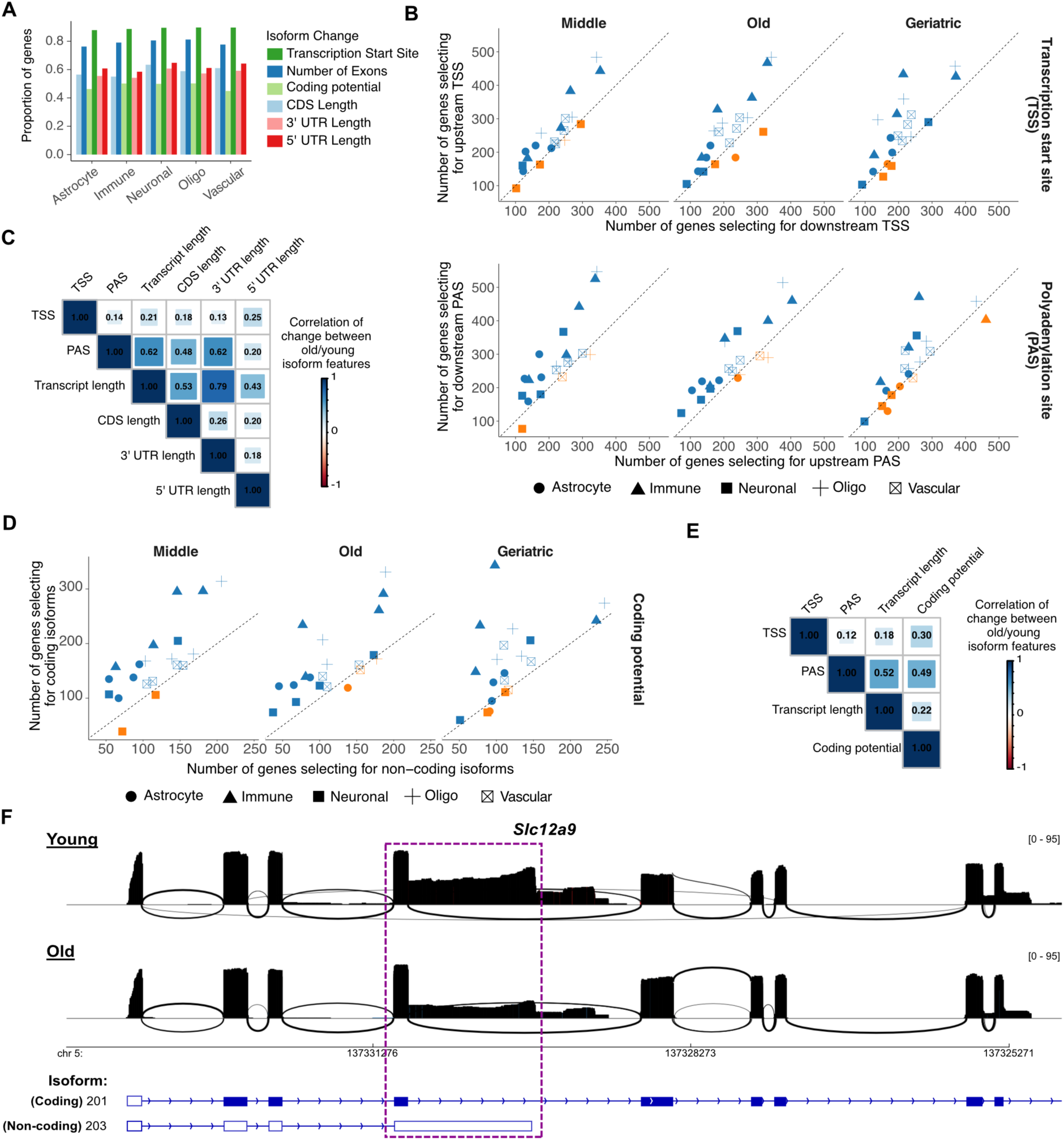
Isoform usage with age associates with upstream transcription and downstream poly(A) site selection. **A)** Bar plot of proportion of genes with significant isoform usage change with age stratified by the type of isoform change. **B)** Scatter plot of the number of genes where isoforms used in young have upstream/downstream TSSs and upstream/downstream PASs, compared to those used in middle, old and geriatric age. For each cell type, the four sex and brain region combinations are shown as independent points. **C)** Spearman’s correlation coefficients of isoform attribute changes for AIGs with at least two protein-coding transcripts. **D)** Scatter plot of the number of genes where isoforms used in young are coding/non-coding compared to those used in middle, old and geriatric age. For each cell type, the four sex and brain region combinations are shown as independent points. **E)** Like (C) but for coding potential and quantified across all AIGs. **F)** Sashimi plots of a partial region of Slc12a9 in the female hippocampus immune cells showing isoform expression in young and old age. Gene models annotated in the Ensembl reference are shown below. Transcript 201 is annotated in Ensembl as coding, whereas 203 as non-coding. The shown gene area has been trimmed at the 3’ end (right) for visualization as transcript 201 was very long. A purple box highlights the area of substantial difference in coverage between young and old.

Given that over 89% of AIGs involved changes to the TSS, we wondered whether selection of downstream TSS defined the observed length and PAS preference. Similar to our analysis for cell-type isoform specificity, we found that TSS change was lowly correlated with isoform length or poly(A) site, indicating that changes in TSS were not quantitatively predictive of absolute isoform length or poly(A) site position (**Fig. 5C**). In contrast, our data again showed correlation between poly(A) site selection and isoform length (ρ = 0.62, **Fig. 5C**) indicating that selection of downstream poly(A) sites resulted in longer transcripts.

Besides changes in TSS and exon number, our results also showed changes in isoform coding potential (**Fig. 5A**). Interestingly, transcriptome wide quantification revealed a global shift in usage in favor of protein coding isoforms with age (**Fig. 5D**), again in contrast to cell-type isoform specificity that had showed the opposite trend. This observation was consistent when comparing young to middle, old and geriatric, again indicating that changes initiate early in adulthood. Comparison of PAS selection with isoform length and coding potential change for all AIGs with at least one non-protein-coding isoform revealed significant correlation (ρ=0.52 and ρ=0.49, respectively, p-val < 10^-256^), indicating that the selection for downstream PAS also associated with selection for longer and coding isoforms (**Fig. 5E**). A characteristic example was the chloride:potassium transporter *Slc12a9* that displayed reduced usage of the shorter, non-coding 203 isoform in favor of the longer coding 201 isoform harboring a downstream poly(A) site in old female hippocampus immune cells (**Fig. 5F)**. Collectively, our data identify hundreds of AIGs that, in contrast to CIGs, show isoform changes that are overwhelmingly driven by upstream TSS selection, downstream PAS, and increase in coding potential.

### Immune cells have increased senescence with age that can be predicted by isoform markers

Increases in inflammatory signaling that accompany aging are increasingly thought to be due to a similar increase in the prevalence of cellular senescence, a state of permanent cell cycle arrest that occurs in response to unresolved DNA damage ^39,40^. Senescent cells contribute to inflammaging through the secretion of senescence-associated factors, including cytokines, chemokines, growth factors, and extracellular matrix proteases, collectively known as the senescence-associated secretory phenotype ^41^. Given that our results showed an increase in immune cells and signaling with age (**Sup. Fig. 13A**), we investigated whether we also saw evidence of senescence in our single-cell data. We developed a senescence score using markers from the SenMayo dataset ^42^ that quantifies the expression of senescence markers for each cell relative to young age and normalized by cell type. Mapping the senescence score across cells revealed a clear, progressive increase in senescence with age that was particularly pronounced in geriatric cells and especially in immune cells (**Fig. 6A**). Quantification of the proportion of senescent cells in each cell type confirmed that immune cells had the greatest number of cells with a high senescence score, consistent with previous studies ^11,43^ (**Fig. 6B, Sup. Table 9**). Notably, senescent cells were more prevalent in the hippocampus compared to the cortex and were especially abundant in females compared to males (**Fig. 6B**).

**Fig. 6:**
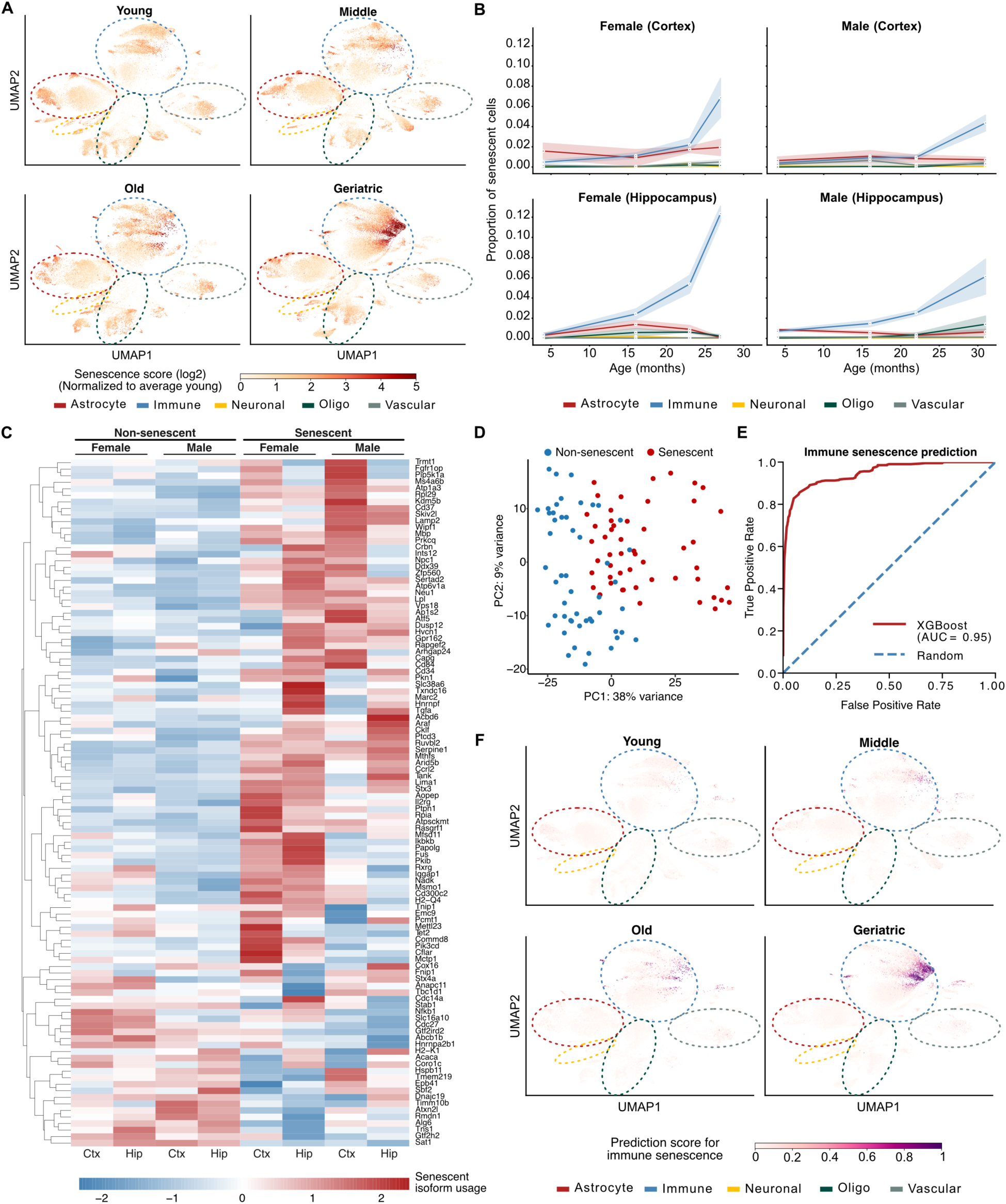
Immune cells have increased senescence with age that can be predicted by isoform markers. **A)** Scatter plots of UMAP isoform expression embeddings overlaid with cell senescence score quantified from senescence markers. Cells have been stratified by sample of origin across young, middle, old, and geriatric age. For reference, dotted ellipses indicate approximate cell-type regions. **B)** Line plots of proportion of senescent cells in each cell-type across aging. **C)** Heatmap of row-normalized isoform usage values of the most highly expressed isoform in senescence. Each row corresponds to a gene identified as having significant differential isoform usage between senescent and non-senescent immune cells. **D)** Scatter plot of the first two isoform usage principal components of isoforms with significantly different usage between senescent and non-senescent cells. **E)** Line plot of true over false positive rate quantified on test set for XGboost model trained on isoform usage values from (C) to classify immune senescent cells. **F)** Same as (A) but overlaid with predicted immune senescence scores from XGboost model.

Discovery of senescence markers has been a major research focus recently, especially with the establishment of the NIH SenNet Consortium ^44^ with efforts concentrating on genes with high expression specificity in senescent cells. We hypothesized that senescent cells may also selectively express unique isoforms that could serve as potential additional markers to gene expression. To investigate this, we compared senescent and non-senescent cells to identify genes with significant isoform usage change, restricting our comparisons to immune cells as they showed the strongest senescence signal. Our analysis identified 105 genes with significant isoform usage changes between senescent and non-senescent immune cells (**Fig. 6C, Sup. Table 10**). Most changes exhibited strong region- and sex-specific patterns, while others were shared across all biological contexts. PCA analysis using isoform usage of the identified genes revealed a clear separation between senescent and non-senescent immune cells in all samples, suggesting that senescence associated isoforms could likely serve as senescence markers in immune cells of the aging brain (**Fig. 6D, Sup. Fig. 13B**).

To test this hypothesis, we developed a machine learning approach using XGBoost to classify senescent over non-senescent immune cells using the isoform usages of the 105 identified genes as input. Quantification of model prediction performance on a left-out test set showed that the model achieved high (0.95) area under receiver operating characteristic curve accurately separating senescent from non-senescent immune cells (**Fig. 6E, Sup. Fig. 13C-D**). To test whether the trained model for immune cells could also classify senescence in non-immune cells we applied it on all other cell types. While predictive performance was substantially lower than immune, our results showed statistically significant separation for all cell types indicating that the model had, at least partially, also learned cell-type agnostic senescent patterns (**Sup. Fig. 13E**).

Finally, to explore how the predicted senescent score could aid in distinguishing and classifying senescent cells, we overlayed the predicted senescence score across the entire dataset on the existing UMAP projection. The updated UMAP plot showed clear separation of senescent cells, particularly evident for geriatric mice (**Fig. 6F**). Overall, our data highlight high rate of acquisition of senescence with aging, particularly in immune cells, and identify novel isoform markers for senescence classification.

## Discussion

The mammalian brain comprises several distinct cell types with unique spatial distributions, functions and aging patterns ^3,6,10^. In this study, we utilized long-read sequencing to examine RNA isoform dynamics in the aging mouse brain at the single-cell level. We constructed a comprehensive dataset across four dimensions with samples from various ages, both sexes, and two brain regions. Since aging is a progressive process, we employed spline modeling to capture both linear and non-linear aging trends starting early in life. To identify sex-specific differences, we analyzed male and female samples separately. Additionally, we measured regional differences by profiling the cortex and hippocampus, two brain areas critical to aging biology.

Our analysis of RNA isoform usage identified notable isoform specificity in distinct cell types across all ages. Our work has shown that alternative TSSs are present in the vast majority of isoform selective events with the selection of the TSS essentially defining isoform usage consistent with previous findings ^33^. Nevertheless, we found that selection of a TSS, upstream or downstream, is not directly associated with the corresponding selection of the PAS or the length of the UTRs. Notably, we found that all cell-types, except for vascular, display specificity towards isoforms that are shorter than the ones used in other cell types, essentially selecting for downstream TSS and upstream PAS. We hypothesize that this preference may be evolutionarily preferred as adaptation of cell-type specific isoforms was likely easier to accomplish within the already optimized sequence space of a gene rather than extending beyond the gene boundaries.

Our results also identified a modest enrichment for non-coding isoforms being cell-type specific that manifested for all cell-types with the notable exception of vascular cells. Given that these non-coding isoforms mostly comprise nonsense-mediated decay substrates expected to degrade, it is intriguing to hypothesize that these isoforms entail a regulatory role controlling RNA abundance in cases of excess energy availability, as is the case in young animals ^17^. Intriguingly, analysis of isoform usage changes with age revealed a distinctly contrasting picture with genes selecting against non-coding isoforms in old age, presumably as a consequence of diminished energy processing and thus availability.

Analysis of aging-related changes in RNA isoform expression also revealed distinct non-linear expression trajectories with time. Interestingly, neuronal RNA isoforms often clustered separately from other cell types highlighting a unique neuronal transcriptome remodeling with age. Notably, we found that neuronal isoforms were the most downregulated with age compared to other cell types. Probing for changes in RNA isoform usage with advancing age revealed hundreds of changes at each life stage. We again found an overwhelming enrichment for changes of the TSS. However, in this case and as with the coding potential, we found that advancing age was associated with a preference for upstream TSS and downstream PAS in stark contrast to cell-type specificity. Notably, neurons showed the highest total isoform usage change in old brains indicating high degree of transcriptomic remodeling with age.

Finally, our data revealed a progressive increase in senescent cell proportion with age in the brain. However, this increase was not uniformly distributed across cell types but instead primarily localized in the immune lineage, consistent with previous studies ^11,43^. Comparative analysis of RNA isoform usage across senescent and normal immune cells identified several isoform markers for cellular senescence. Our results showed that these markers have sex and brain region specificity with a subset showing consistent change across all biological modalities. Discovery of senescence markers has been the subject of intense study with the establishment of an NIH initiative and consortium to map senescent cells ^44^ and we thus anticipate RNA isoform diversity will add an additional layer for senescence characterization. As proof of principle, we developed a machine learning approach to classify senescent from non-senescent cells based on isoform usage. Our results showed high prediction accuracy indicating RNA isoform diversity as an additional layer of the senescence phenotype. In conclusion, our work establishes an RNA isoform atlas of the adult and aging mouse brain and discovers RNA isoform selection dynamics that define transcriptomic state and cell function in aging.

## Methods

### Animals

All animal procedures were approved by the Animal Care and Use Committee of the National Institute on Aging (Animal Study Protocol: 483-LGG-2025). Male and female C57BL/6JN mice were used in this study. Mice were provided a standard diet (Teklad Global 18% Protein Rodent Diet, Envigo, Indianapolis, IN, USA) and maintained on a 12 h light/dark cycle at 20-22 C. Young (4 months; n=3 males, n=3 females), middle age (16 months; n=3 males, n=3 females), old (22, 23 months; n=3 males, n=3 females) and geriatric (31 months; n = 5 males, 27 months; n = 3 females).

### Tissue dissociation

Brains were extracted and immediately placed on an ice-cold petri dish for dissection of the prefrontal cortex (1/2 hemisphere) and hippocampus (1 hemisphere), which were processed separately for the remainder of the study. Adult mouse brain dissociation was performed using the Miltenyi Adult Brain Dissociation Kit (Mouse and rat; Miltenyi Biotec, Auburn, CA; Cat. #130-107-677). All steps were performed on wet ice unless otherwise noted. Dissected regions were transferred to 15 ml falcon tubes containing cold Dulbecco’s PBS (D-PBS; Thermo Fisher Scientific; Asheville, NC; Cat. #14040133) and gently inverted 5x to remove blood and extraneous debris. Tissue was transferred to gentleMACS C Tubes (Miltenyi Biotec, Auburn, CA; Cat. #130-096-334) containing 1950 µl Enzyme Mix 1 (50 µl Enzyme P + 1900 µl Buffer Z; Miltenyi Biotec, Auburn, CA), followed by addition of 30 µl Enzyme Mix 2 (20 µl Buffer Y + 10 µl Enzyme A; Miltenyi Biotec, Auburn, CA). Tubes were placed on a gentleMACS Octo Dissociator with Heaters (Miltenyi Biotec, Auburn, CA; Cat. #130-096-427) and the appropriate gentleMACS program was run for given tissue input weight (37C_ABDK_02). Following termination of the program, C-Tubes containing tissue were detached and briefly centrifuged at 300 *g* at 4°C to collect dissociated tissue to the bottom of the tube. Sample was gently resuspended 10x using a 10ml serological pipette and applied to a pre-moistened 70 µM MACS SmartStrainer (Miltenyi Biotec, Auburn, CA; Cat. #130-098462) on a 50m-l tube. Next, 10 ml of cold D-PBS was added to the C Tube which was then closed, gently inverted 10x, and applied to the same 70µM MACS SmartStrainer to ensure collection of all remaining sample. C Tube and MACS SmartStrainer were discarded, and 50 ml tube containing tissue and D-PBS was centrifuged at 300 *g* for 10 minutes at 4°C, after which supernatant was aspirated completely.

Cell pellet was resuspended in 1550µl D-PBS and transferred to a 5ml Eppendorf tube, followed by addition of 450 µl of Debris Removal Solution (Miltenyi Biotec, Auburn, CA) and gentle mixing using a 1000µl pipette tip. Cold D-PBS (2ml) was gently overlayed on top of cell suspension and debris removal solution mixture, so as not to disrupt phases. Sample was centrifuged at 300 *g* for 10 minutes at 4°C and halted with natural deceleration and no brake, so as not to disturb the pellet. The top two phases of mixture were removed from the suspension and 2 ml fresh, cold D-PBS was added to the tube which was then gently inverted 3x. Sample was centrifuged at 1000 *g* for 10 minutes at 4°C with full acceleration and full brake, after which supernatant was aspirated completely.

To remove extraneous red blood cells, Red Blood Cell Removal Solution (10X, Miltenyi Biotec, Auburn, CA) was diluted 1:10 with double-distilled water, and 0.5 ml was added to the cell pellet, resuspended, and incubated at 4°C for 10 minutes. An appropriate amount of cold PB buffer (D-PBS + 0.5% bovine serum albumin) was added to the tube then centrifuged at 300 *g* for 10 minutes at 4°C. Supernatant was aspirated completely, and cell pellet was resuspended in D-PBS without calcium, magnesium (ThermoFisher Scientific; Asheville, NC; Cat. #14190144) / 0.04% BSA (Miltenyi Biotec, Auburn, CA; Cat. #130-091-376) / 0.1% Protector RNase Inhibitor (Sigma Aldrich, St. Louis, MO; Cat# 3335402001). Cell suspension was transferred to a 2 ml DNA-LoBind tube on ice (Eppendorf, Westbury, NY; Cat. #022431048) to ensure accurate counting.

To quantify cell viability, 18 µl of cell suspension was mixed with 2 µl Propidium Iodide/Acridine Orange (Logos Biosystems; Annandale, VA; Cat. #F23011) in a 2 ml LoBind tube. 10 µl was then loaded to each side of an Ultra-low Fluorescence Photon Slide (Logos Biosystems; Annandale, VA; Cat. #L12005) and inserted into a LUNA-FL Dual Fluorescence Cell Counter (Logos Biosystems; Annandale, VA; Cat. #L20001) for viewing and counting. If excess debris remained, cell suspension was filtered through a 70 µM FlowMi Strainer (Bel-Art; Pequannock, NJ; Cat. #H13680-0070) and an additional aliquot of 18 µl was removed for recounting. Cell suspension was diluted to a final concentration of approximately 1000 live cells/µl using D-PBS/0.04% BSA/0.1% Protector RNase Inhibitor before proceeding with the 10x Genomics protocol.

### Single-cell, long-read sequencing library preparation

Single-cell cDNA was generated using the Chromium Next GEM Single Cell 3’ GEM, Library & Gel Bead Kit v3.1 (Cat #1000128, 10x Genomics) following the manufacturer’s protocol (version CG000204 Rev D). Single cell suspensions from cortex or hippocampus were mixed with Gel Beads to target the recovery of 6000 cells. Single-cell Gel Beads-in-Emulsion (GEMs) were generated on the 10x Chromium Controller. Reverse transcription of barcoded RNA was performed in each GEM and during this process, each cell was assigned a unique barcode, and each RNA transcript was labeled with a unique molecular identifier (UMI). The barcoded cDNA was amplified by PCR for a total of 12 cycles, such that it yielded sufficient material for library preparation while minimizing the amplification artifacts. The amplification products were cleaned with SPRI beads and were subsequently used for long-read library preparation. 10 ng of single-cell barcoded cDNA were used to prepare long read cDNA-PCR libraries following the manufacturer’s protocol (version SST_v9148_v111_revB_12Jan2022, SQK-PCS111, Oxford Nanopore Technologies) except for the post-pull-down PCR step that was realized with 6 cycles instead of 4. Final libraries were quantified using Qubit 1X dsDNA High Sensitivity (HS) assay kit (ThermoFisher Scientific, Q33231) and analyzed on a BioAnalyzer (High Sensitivity DNA kit, Agilent, 5067-4626). Libraries were loaded to R9.4.1 flow cells (ONT, FLO-PRO002) and run on a PromethION device (ONT, PRO-SEQ024).

### Long-read sequencing data processing

Long-read sequencing data preprocessing was accomplished by using the ONT Sockeye (v0.4.0) pipeline with custom workflow modifications. Briefly, in the Sockeye pipeline basecalling of nanopore data was done using the ONT Guppy (v.6.1.2) basecaller. After basecalling, read quality control and adapter identification was performed using custom Python scripts which searched for adapter pairs. Alignment to the mm10 reference genome was performed using the minimap2 (v.2.24) aligner. Following alignment, barcode identification and filtering of aligned reads was performed using SAMtools (v.1.14) and Pysam. Once reads with 10x genomics barcodes were identified, reads with the same UMI entry were collapsed, and a collapsed BAM file for each library was created. Gene-level identification was then performed using sockeye scripts, which produced barcode-gene matrixes for each sequencing library.

Isoform-level data was processed using Stringtie2 (v2.2.1) and quantified on the Ensembl annotation (v98). From the UMI-collapsed BAM file, Stringtie2 was used to build GTF files of the transcriptomes for each sample. All GTF files were subsequently merged into a consensus transcriptome for all libraries. The UMI collapsed BAM files were then realigned to the consensus transcriptome. Only reads aligning to Ensembl annotated transcripts were kept for further analysis. These alignment files were used to create an isoform-barcode matrix using custom python scripts.

### Single-cell cluster identification and annotation

Barcode-gene assignments were processed with the scanpy (v.1.9.3) library ^45^. For quality control, barcodes with fewer than 100 distinctive assigned genes or more than 20% mitochondrial content were removed. In addition, barcodes with more genes than the 98% of barcodes were also removed from analysis. After barcode filtering, samples were integrated using the scVI (v.0.20.3) toolkit ^46^ as previously suggested. Computational clustering was performed using the Leiden algorithm ^47^ with default parameters. A panel of established marker genes from the PanglaoDB and Tabula Muris dataset was used to assign Leiden cell clusters to known cell types. The same cell-type annotations were subsequently used for cell-isoform counts.

### Transcript filtering and pseudobulk quantification

To assure results were completed with the most well-annotated transcripts, only transcripts with high (TSL1) support level from the Ensembl database were used. Furthermore, transcripts that appeared in fewer than 0.5% of the cells of a given cell type in a sex-region combination were removed from the analysis. Subsequently, cell-type gene and isoform expression quantification were performed by pseudobulk analysis (**Sup. Table 11**). Filtered transcripts were normalized using DESeq2 at the cell and sample level (v1.42.0) ^48^.

### Cell-type isoform selectivity

To identify isoform-selective genes, isoform usage was quantified as the ratio of counts of a given isoform over all isoform counts of the gene. To identify isoforms specifically selected in a cell type of interest, the isoform usage values for a given cell type were compared against the usage values for all other cell types using DRIMSeq ^31^. and statistically significant changes were identified. The feature-level option to extract effect size and p-values associated with each isoform was enabled. Isoforms with statistically significant lower (p<0.05) usage in the cell type of interest compared to the other cell types were considered “low usage”, while those with higher usage were considered “high usage”.

### Isoform usage-weighted attribute analysis

To retrieve a quantifiable measure of the difference in the usage of individual isoform attributes in a CIG or AIG we devised a new metric that weights the attribute of interest by the corresponding isoform usage in a gene. Specifically, given a gene *g* with isoforms *g*_*i*_, *i* = 1.. *N* each with usage *u*_*i*_ and value of the attribute of interest *a*_*i*_ then the weighted-attribute metric *A*_*g*_ is calculated as

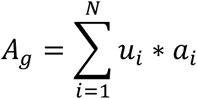

While this calculation is straightforward for isoform length, both
poly(A) and TSS genomic coordinates required normalization. To accomplish that we normalized TSS and poly(A) sites relative to the most upstream site amongst the corresponding gene isoforms. The most upstream site was set at 0, and all other sites were quantified by their binned distance from that site. The bin size was selected at 20 nts for TSSs and 75 nts for poly(A) sites similar to ^33^. Subsequently, the relative binned distance was used as the value *a*_*i*_. Similarly for coding potential, protein-coding isoforms were encoded as 1 and non-coding isoform as -1.

When calculating the protein-coding attributes of UTR and CDS lengths, only protein-coding genes with at least two protein coding isoforms were considered, and transcripts with unclear or ambiguous annotation were removed.

When comparing two conditions, either one cell-type against all others or one-age against all others, we quantified the difference between the weighted-attribute averages. To maintain consistency between the poly(A) site and TSS comparisons, the TSS difference between conditions was negated so that a positive value would be associated with change towards upstream TSSs. Overall, a positive change corresponded to longer isoform, CDS, 5ʹ UTR and 3ʹ UTR length, upstream TSS, downstream poly(A) site, and higher protein coding potential.

### Identification of isoform expression clusters with age

Using the filtered pseudobulk transcripts, isoform expression across aging was further investigated by grouping statistically significantly changing isoform expressions into distinct clusters. This was accomplished using two steps. The first was to determine if an isoform-cell expression was significantly changing with time. To capture non-linear patterns, a 2^nd^ order polynomial function was fitted to each expression and the null hypothesis was constant expression with age. A likelihood ratio test was then used to compare the null hypothesis function and the fitted function, and a threshold of p<0.01 was used to filter transcripts. After obtaining the filtered isoform-cell expressions, they were clustered together by the biological modalities of sex and brain region using the TMixClust (v1.22) package ^49^. The TMixClust package uses a Gaussian mixed-effects model to group each isoform-cell expression into clusters using each contig’s mean and variance.

### Isoform length change with age

To evaluate systemic isoform length changes first a pairwise differential transcript expression between young age and all other time points was performed using DESeq2. To remove noisy transcripts, we kept transcripts whose expression was in the top 50% in either the young age, or the older age time point. Transcripts were subsequently binned into three groups based on increasing transcript length. The distribution of the log2 fold change was then compared between each of these groups using a pairwise Wilcoxon test.

### Isoform usage change with age

To quantify isoform usage changes with age, a pairwise analysis of isoform usage using DRIMSeq was performed. All aging timepoints were compared and a statistical significance threshold (p < 0.05) was used to determine if there was an isoform usage change with age.

Isoform attribute change analysis was performed as for cell type specificity by quantifying the isoform usage-weighted attribute metric. Pairwise aging comparisons were performed between young/middle, young/old and young/geriatric, selecting the usage values from only those time points.

### Senescence score quantification

A cell senescence score was calculated by normalizing the raw total counts of each cell to 10,000 and subsequently calculating the mean expression of all isoforms from the top 20 murine gene markers listed in Table 2 in the SenMayo dataset ^42^. To account for differences in marker expression in different cell types, a normalized score was calculated by dividing the senescence score of each cell by the average senescence score of the young cells of the corresponding cell type. Cells whose normalized senescence score was over 5, selected to approximate the background distribution observed in young cells, were defined as senescent and those that were below as non-senescent.

After quantifying the proportion of senescent cells per cell type across ages, a distinctive aging-related pattern emerged in immune cells thus for subsequent analysis only the immune senescent and non-senescent cells were used. Differential isoform usage analysis between senescent and non-senescent cells was performed using DRIMSeq by incorporating all time points. Genes with significant isoform usage change (padj. < 0.01) between senescent and non-senescent cells were selected as markers of senescence (**Sup. Table 12, 13**).

### Machine learning for isoform markers of immune senescence

A senescence classification machine learning model was developed using cells of the immune lineage, which was found to contain the highest proportion of senescent cells. Isoforms with statistically significant usage changes (Dirichlet-multinomial model, p < 0.01) between senescent and non-senescent immune cells were selected as features, resulting in 741 isoforms from 105 genes. Binary classification labels were created based on the normalized senescence score: cells with a score below 4 were labeled as non-senescent (label 0), and cells with a score above 7 were labeled as senescent (label 1). The resulting dataset, which comprised 53,120 non-senescent and 4,125 senescent cells, was split into training (90%), validation (5%), and test (5%) sets, with balanced representation across label, age category, sex, and brain region. XGBoost was employed as the classification model, and 5-fold cross-validation was used for hyperparameter tuning. In the cross-validation, a learning rate of 0.3, 10,000 boosting rounds, and 50 early stopping rounds were utilized. The final model was trained with the following parameters: a learning rate of 0.01, binary logistic objective, area under the precision-recall curve as the evaluation metric, approximate tree method, max depth of 7, minimum child weight of 16, subsample of 0.9967762267267729, subsample ratio of columns for each split of 0.4100293732438932, L2 regularization term on weights of 6.001086886655973, 10,000 boosting rounds, and 50 early stopping rounds. Default values were used for all remaining parameters.

## Acknowledgements

This work utilized the computational resources of the NIH HPC Biowulf cluster (http://hpc.nih.gov).

## Author contributions

MM and MRC conceived the project. AR performed computational analysis with help from KG, MJP, SAD, and CTL. MD performed tissue dissociation. NY performed single-cell preparation. LW performed tissue dissection with help from SM. CB performed nanopore library preparation. AR and MM interpreted the results assisted by MRC and PS. AR and MM wrote the manuscript with feedback from all authors.

