## Supplemental Figures for "Cell-specific RNA isoform remodeling in the aging mouse brain"

**Supplementary Figure 1**

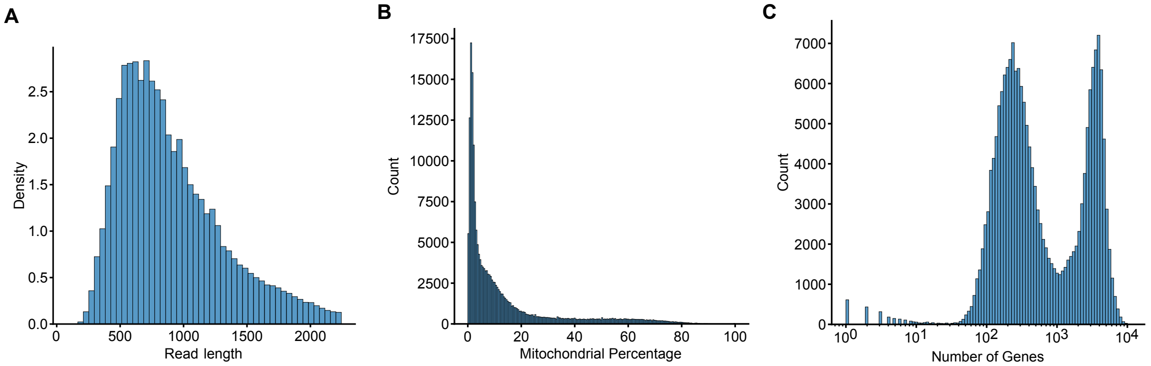


**Sup. Fig. 1: Single-cell long-read sequencing statistics. A-C)** Histogram of read length from a random sample (5%) of all sequenced reads (A), of the mitochondrial content in cells (B) and number of distinct genes per cell barcode (C).

**Supplementary Figure 2**

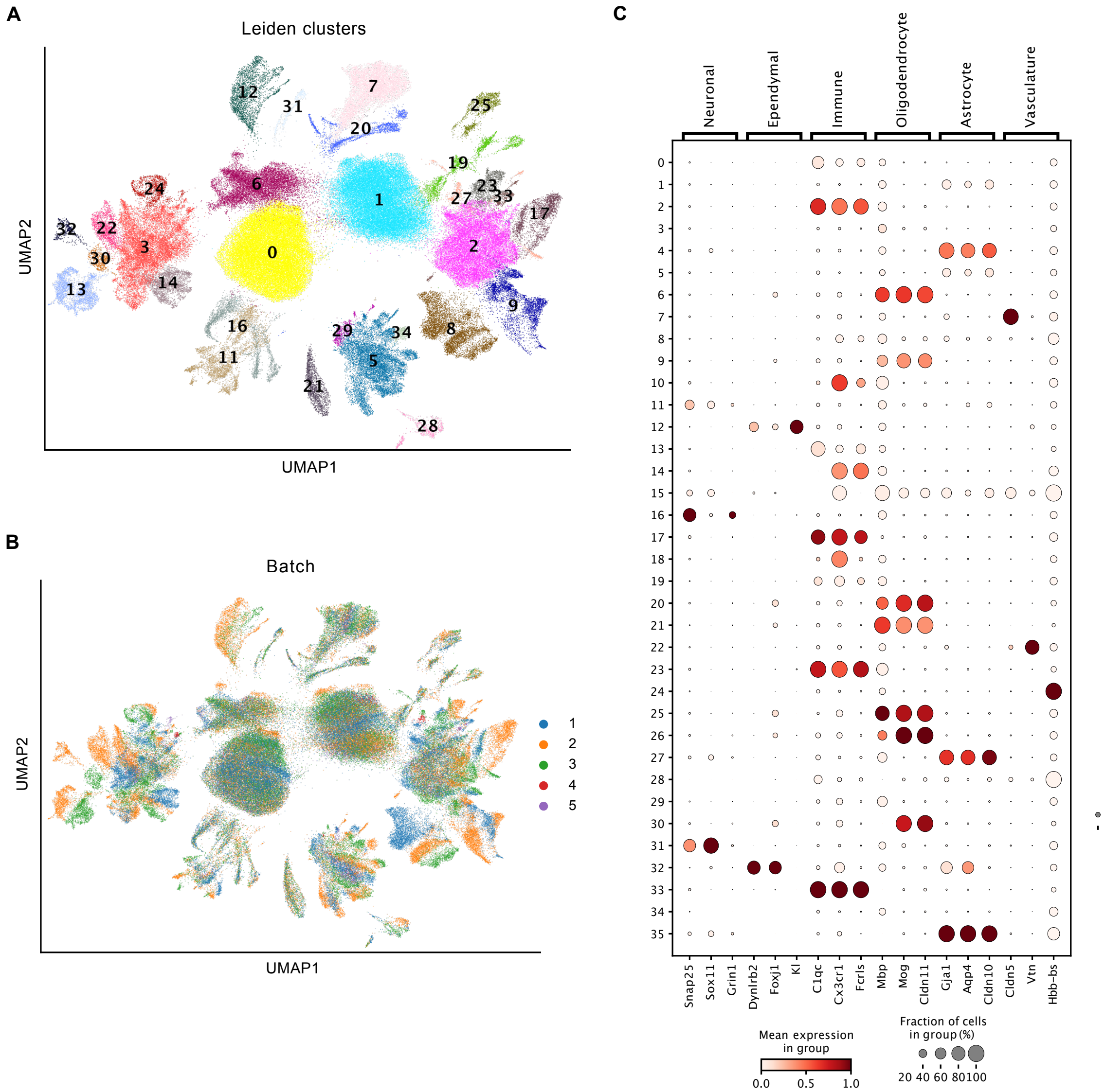


**Sup. Fig. 2: Cell type annotation based on gene markers. A-B)** Scatter plot of UMAP gene expression embeddings with computationally identified Leiden clusters and sequencing batches colored. **C)** Dot plot of average marker gene expression in Leiden clusters.

### Supplementary Figure 3

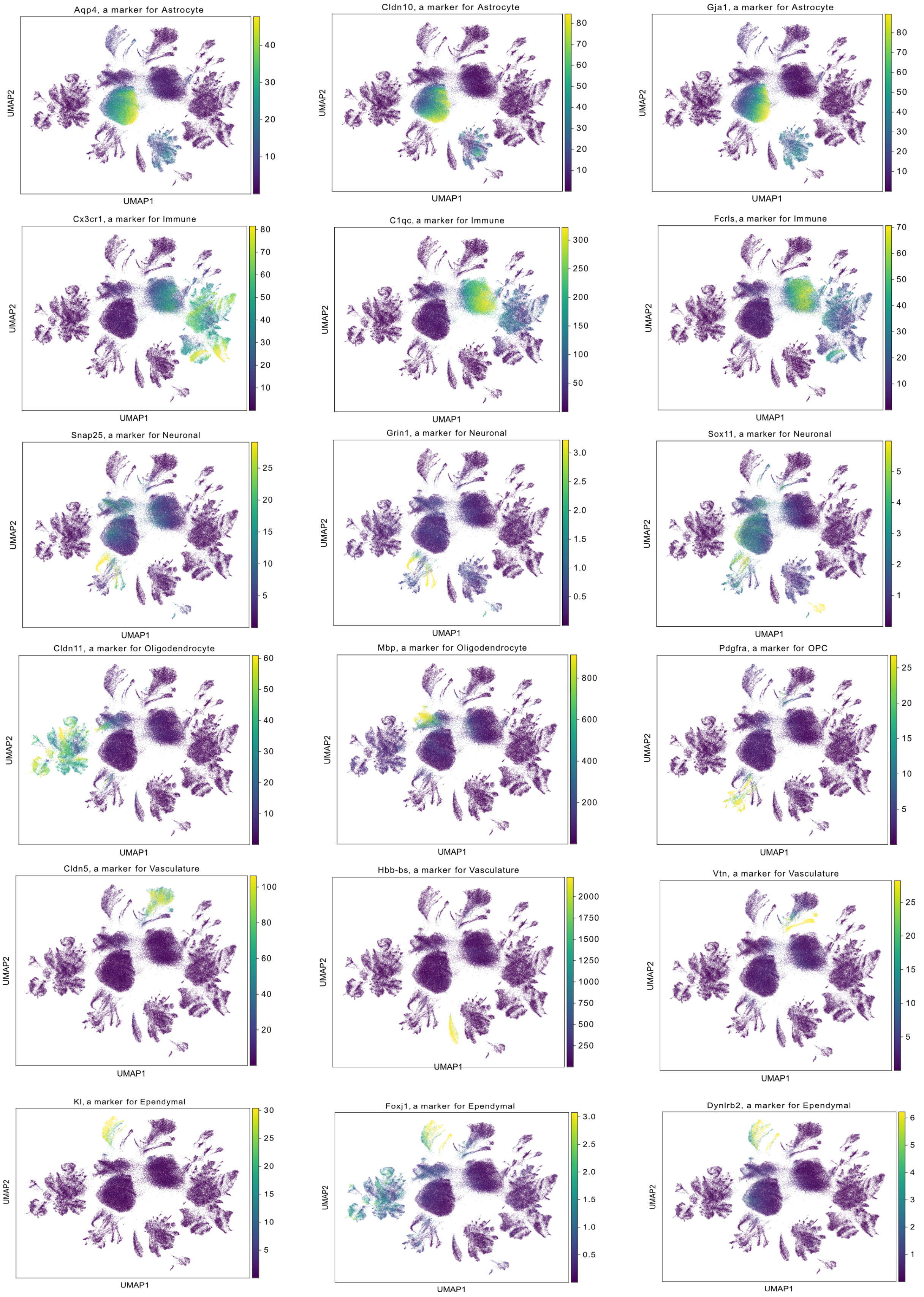

**Sup. Fig. 3: Marker gene expression in cells. A)** Scatter plots of UMAP gene expression embeddings colored by the expression of noted marker genes.

**Supplementary Figure 4**

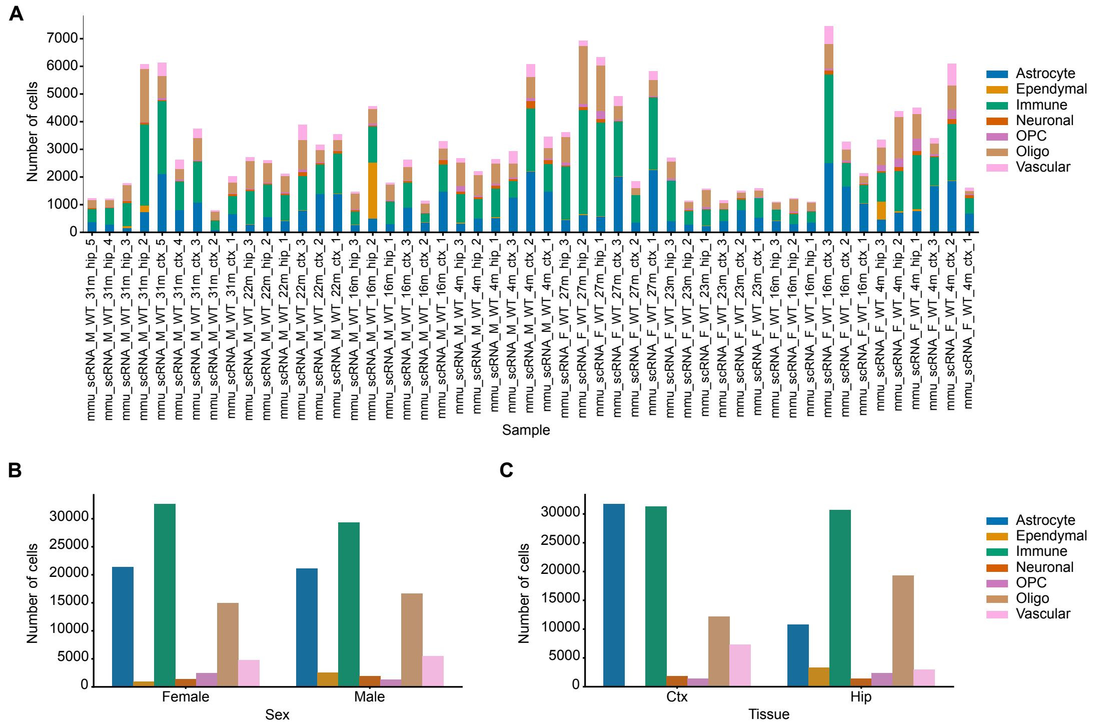


**Sup. Fig. 4: Cell counts distribution per sample, sex, and brain region. A)** Bar plot of cell type composition in each sample. **B-C)** Bar plot of cell type composition for sex (B) and brain region (C).

**Supplementary Figure 5**

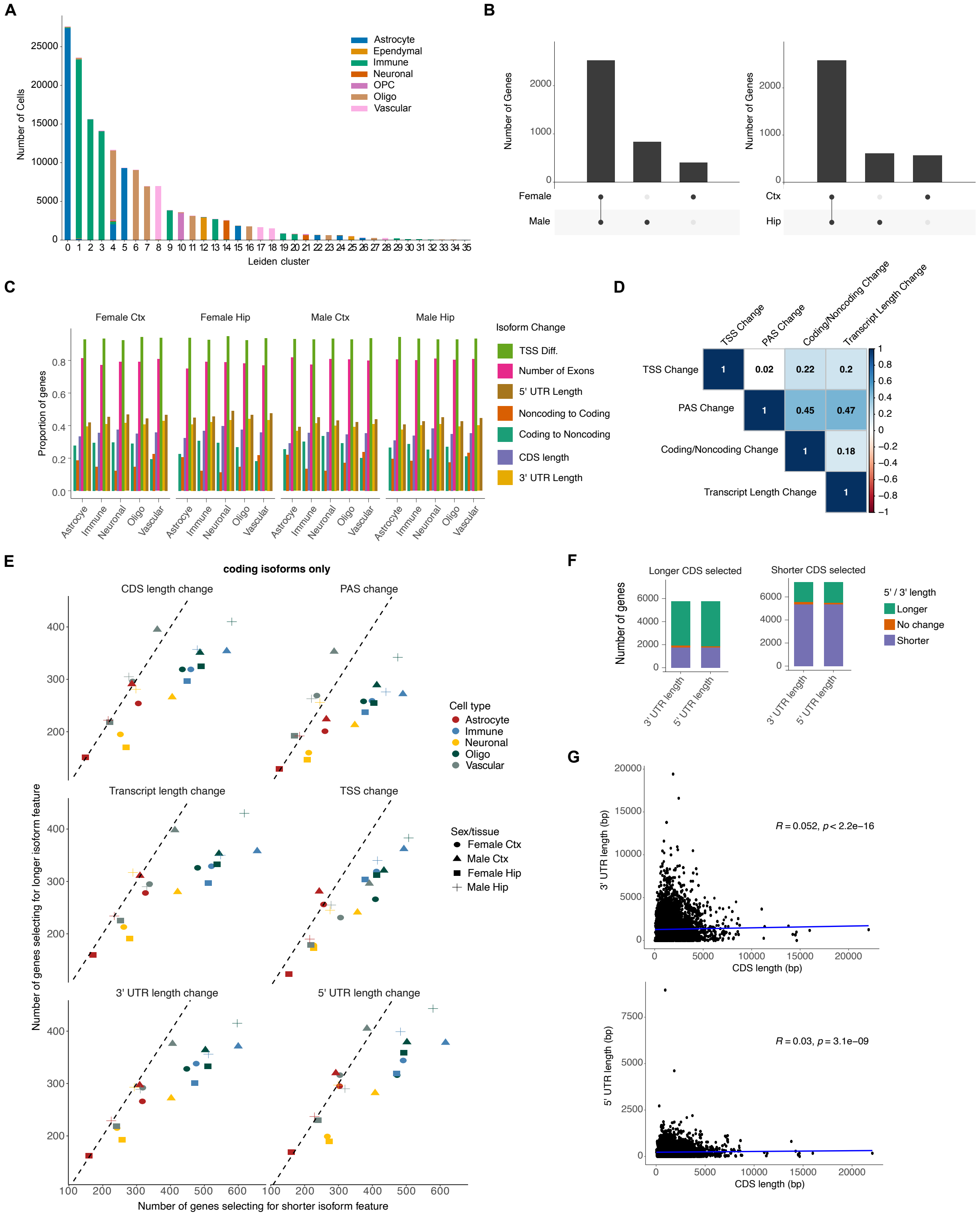


**Sup. Fig. 5: Features of isoforms identified as cell-type specific. A)** Bar plot of cell-type composition, defined from gene abundance, in Leiden clusters constructed from isoform abundance. **B)** Upset plots quantifying the overlap between sexes and brain regions of genes identified as having cell-type isoform specificity. **C)** Bar plot of proportion of significant isoform-specific genes stratified by the type of change between the cell-specific and non-specific isoform stratified by sex and brain region. **D)** Spearman correlations between changing features of cell-type-specific and non-specific isoforms for genes with at least one non-coding isoform. **E)** Scatter plot of isoform changes of protein coding isoforms in genes with at least two protein coding isoforms. Shown are changes in CDS length, poly(A) site selection, transcript length, TSS, 5' UTR length, and 3' UTR length. **F)** Bar plots of number of genes identified having significant cell-type isoform specificity genes and with longer or shorter 5' and 3' UTRs stratified by the corresponding CDS length change. **G)** Scatter plots of CDS and 3' UTR length (top) or 5' UTR length (bottom) for all transcripts annotated in the Ensembl database.

**Supplementary Figure 6**

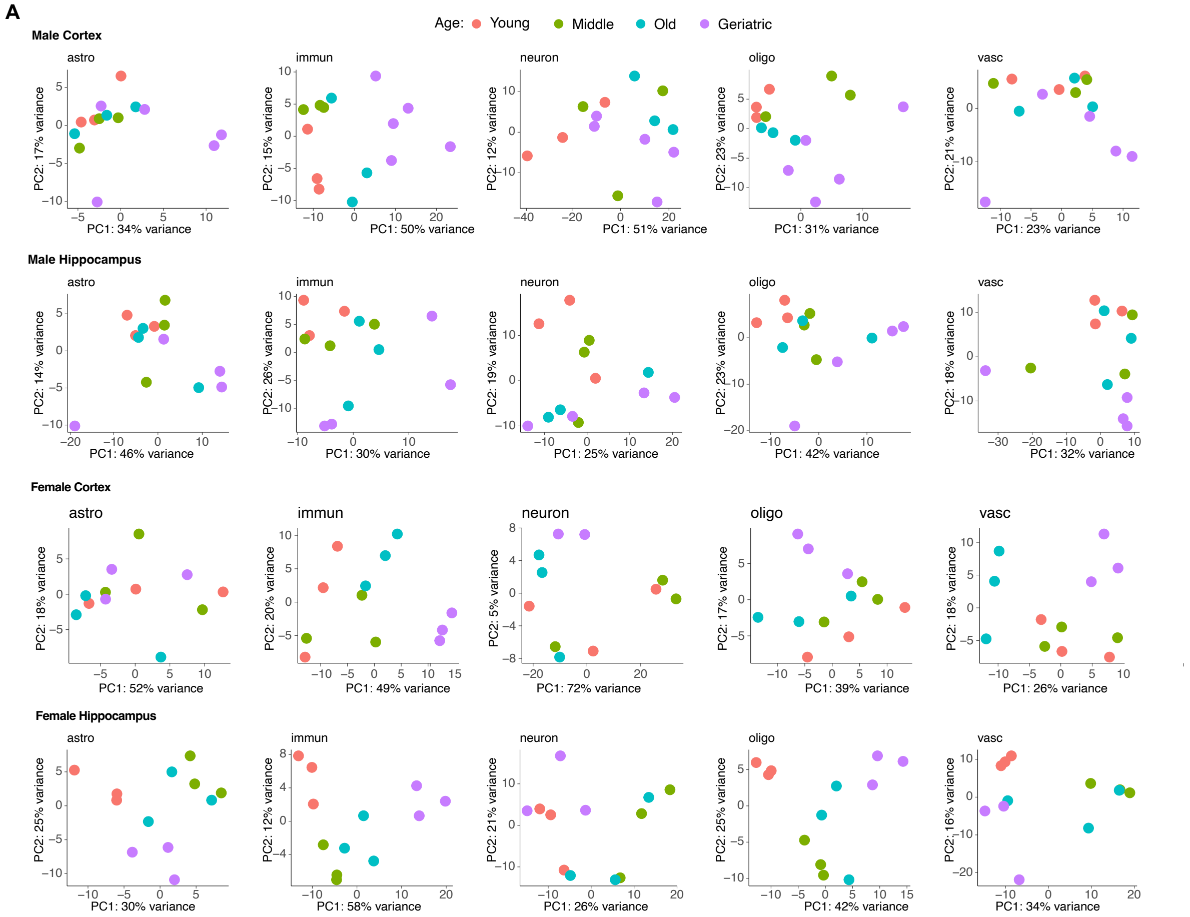


**Sup. Fig. 6: Principal component analysis of all samples with aging. A)** Scatter plots of PCA of isoform expression of all samples stratified by sex, brain region and cell type and colored by age.

**Supplementary Figure 7**

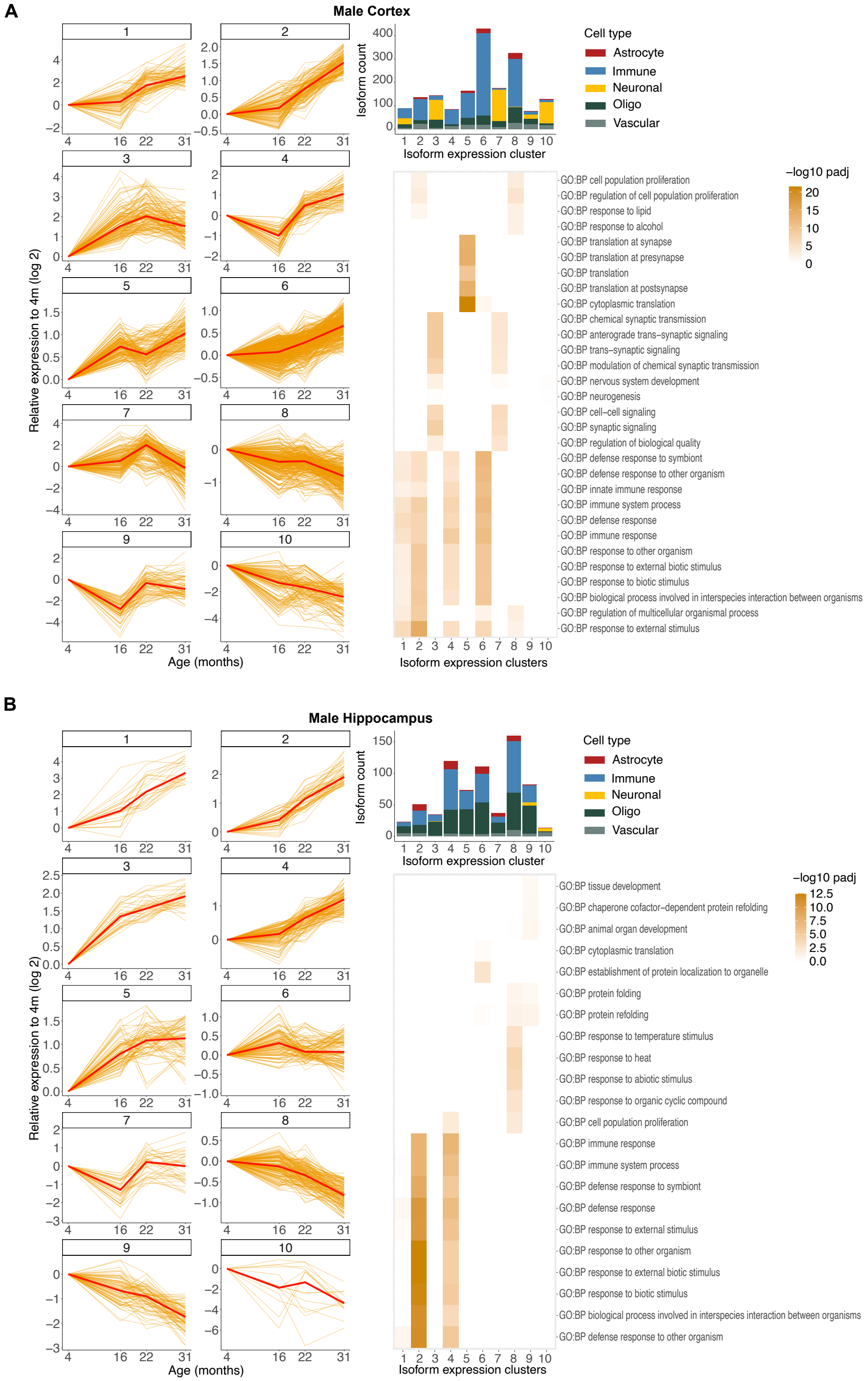


**Sup. Fig. 7: Male isoform expression trajectories with age. A-B)** Line plots (left) of isoform expression at different ages in the male cortex (A) and hippocampus (B) clustered by expression trajectory. Each line represents isoform expression within a cell type, normalized to the corresponding expression at 4 months. The dark red line corresponds to the average expression of the cluster. Bar plot (top right) of cellular composition of each isoform expression cluster.

Heatmap (bottom right) of significance of gene ontology terms enrichment in each cluster. Enrichment is calculated against all expressed genes in the male cortex and male hippocampus, respectively.

**Supplementary Figure 8**

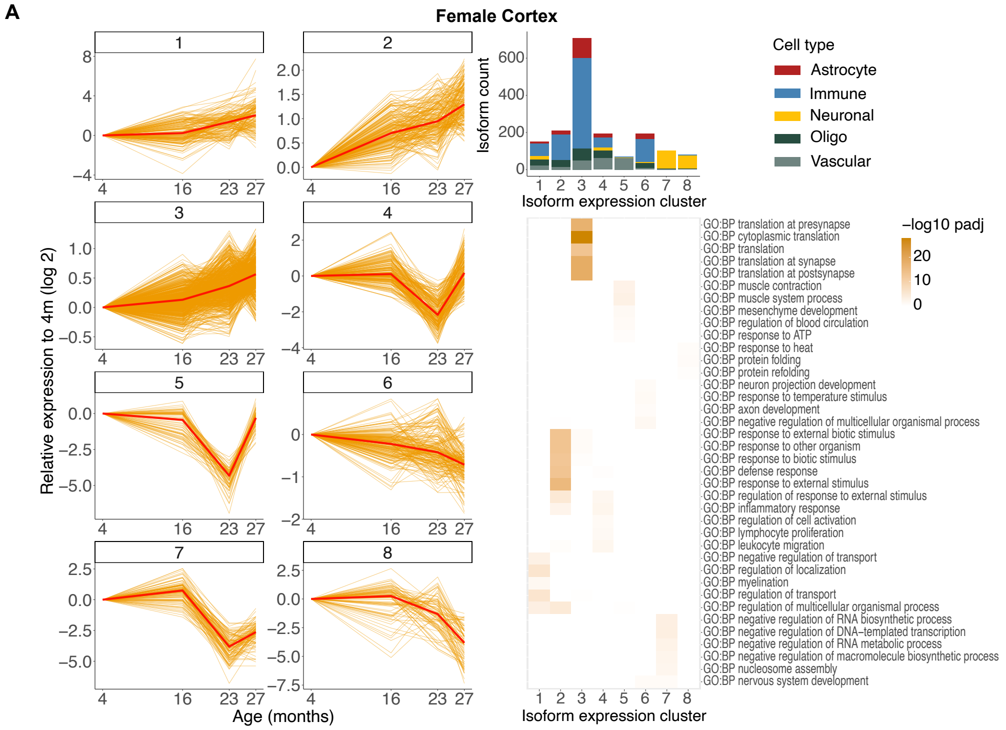


**Sup. Fig. 8: Female isoform expression trajectories with age. A)** Line plots (left) of isoform expression at different ages in the female cortex clustered by expression trajectory. Each line represents isoform expression within a cell type, normalized to the corresponding expression at 4 months. The dark red line corresponds to the average expression of the cluster. Bar plot (top right) of cellular composition of each isoform expression cluster. Heatmap (bottom right) of significance of gene ontology terms enrichment in each cluster. Enrichment is calculated against all expressed genes in the female cortex.

### Supplementary Figure 9

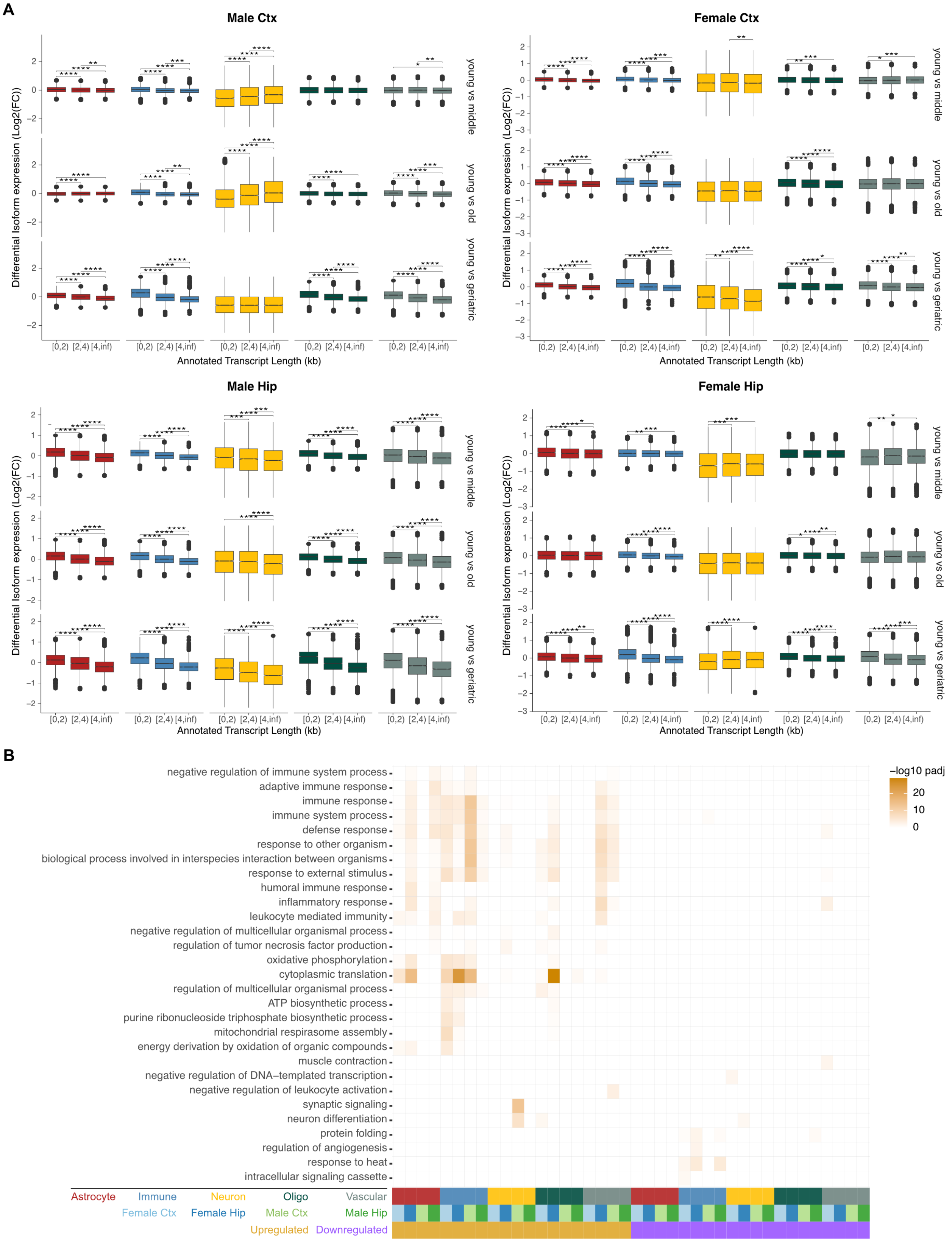

**Sup. Fig. 9: Age-associated isoform length imbalance with age and GO analysis. A)** Box plots of isoform expression change for isoforms with significant expression change between indicated ages stratified by average annotated isoform length, sex and brain region. **B)** Heatmap of enrichment p-value of the most statistically significant GO terms for isoforms upregulated or downregulated with age, stratified by cell type, sex and tissue. Enrichment is calculated against all expressed genes in the corresponding sex, brain region and cell-type.

**Supplementary Figure 10**

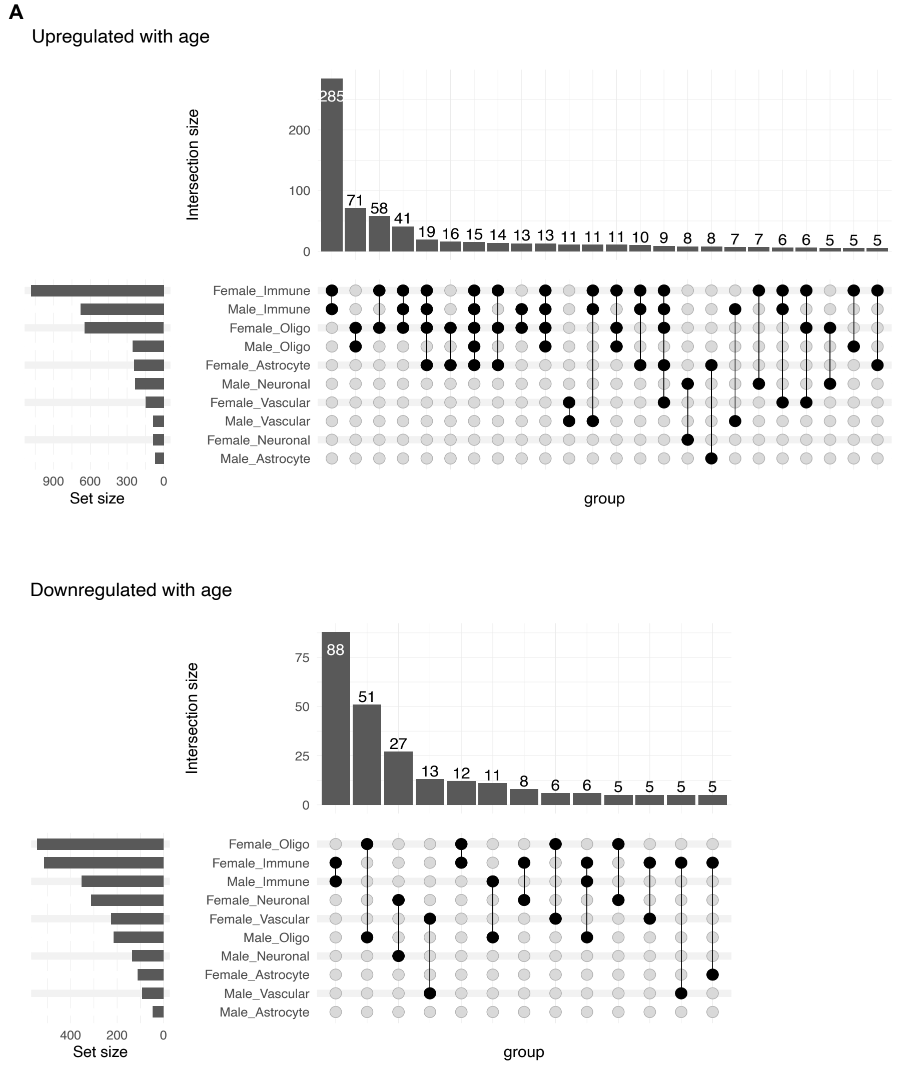


**Sup. Fig. 10: Overlap of isoform expression changes between cell type and sex. A)** Upset plots of the number isoforms with significant aging-associated expression change that overlap between sex and cell types stratified by upregulated and downregulated with age.

**Supplementary Figure 11**

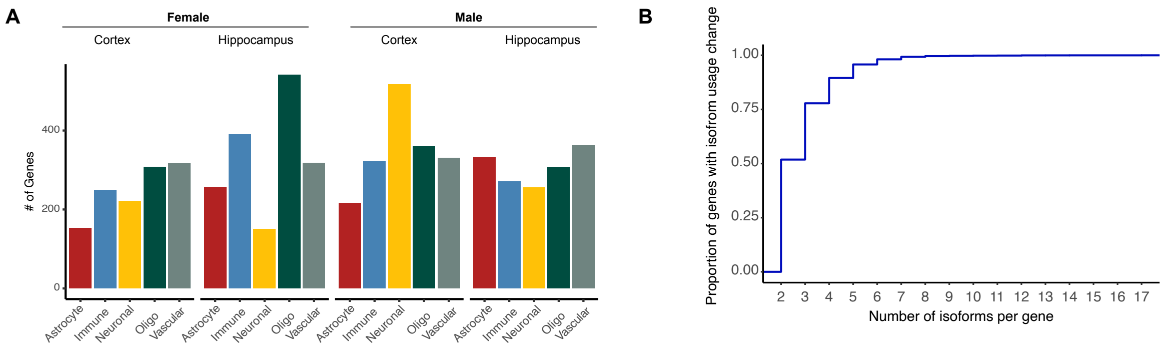


**Sup. Fig. 11: Characterization of genes with age-related differential isoform usage. A)** Bar plot of number of genes with differential isoform usage with aging across sex, brain region and cell type. **B)** Empirical cumulative distribution plot of gene count versus the number of isoforms within the gene that are changing usage.

**Supplementary Figure 12**

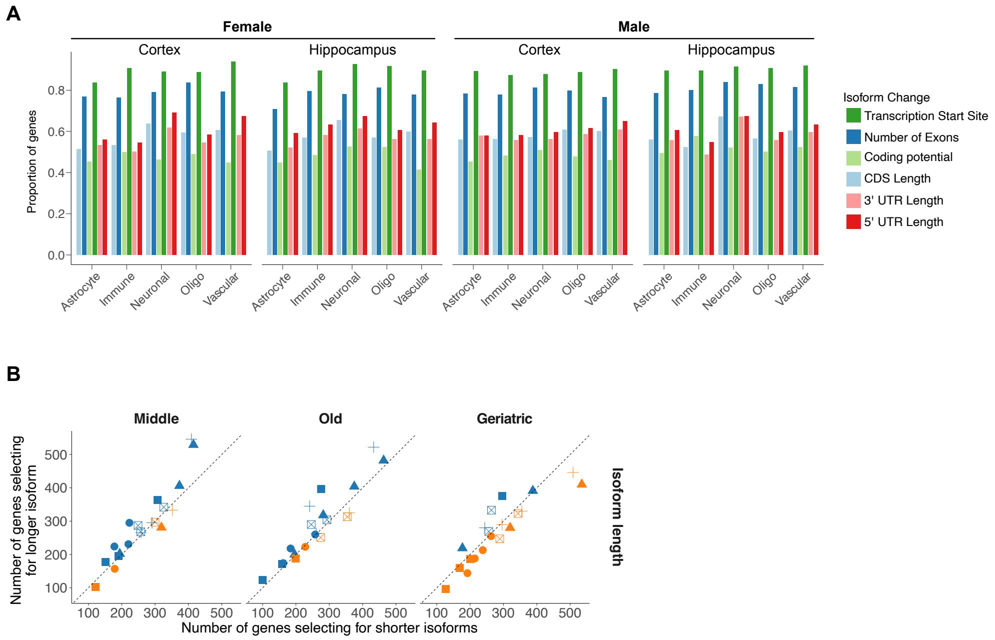


**Sup. Fig. 12: Features of isoforms identified as changing usage with age. A)** Bar plot of proportion of genes with significant isoform usage change with age stratified by the type of isoform change per sex and brain region. **B)** Scatter plot of the number of genes where isoforms used in young are longer/shorter compared to middle, old and geriatric age. For each cell type, the four sex and brain region combinations are shown as independent points.

**Supplementary Figure 13**

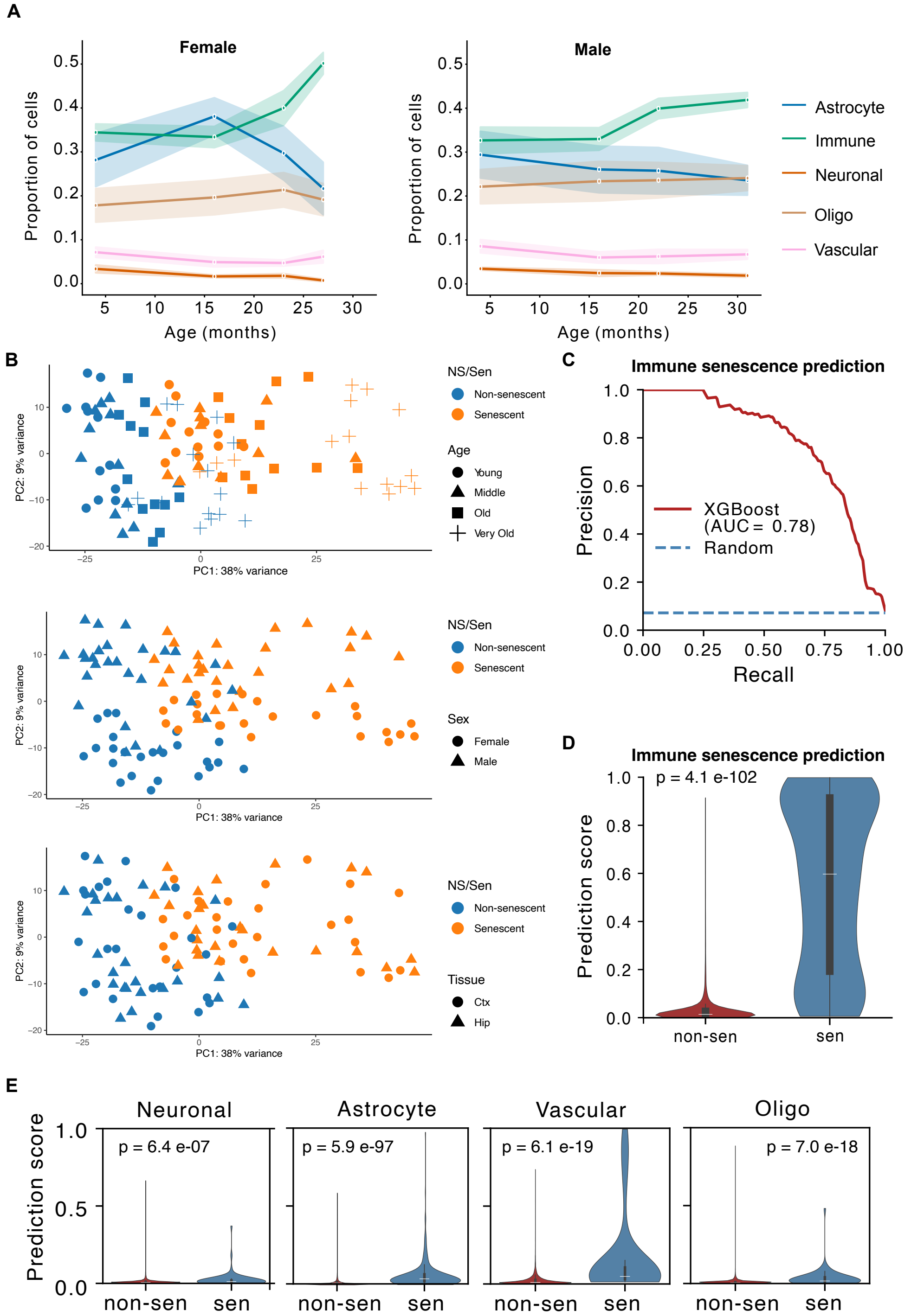


**Sup. Fig. 13: Immune cells have increased senescence with age that can be predicted by isoform markers. A)** Line plot of average cell proportion in each sample for females and males. **B)** Scatter plots of PCA showing data separation along senescence/non-senescence axis and aging (top), sex (middle), and tissue (bottom). **C)** Line plot of precision-recall curve quantified on test set for XGboost model trained on isoform usage values to classify immune senescent cells. **D)** Violin plot of senescence prediction score for senescent and non-senescent immune cells quantified from XGboost model trained on immune cells. The Mann-Whitney p-value is also shown. **E)** Same as (D) but for senescence prediction on neuronal, astrocyte, vascular and oligodendrocyte cells. The XGboost model trained on immune cells was used for prediction.
